# Wnt/beta-catenin signalling is dispensable for adult neural stem cell homeostasis and activation

**DOI:** 10.1101/2020.12.29.424694

**Authors:** Sophie H. L. Austin, Lachlan Harris, Oana Paun, Piero Rigo, François Guillemot, Noelia Urbán

## Abstract

Adult mouse hippocampal neural stem cells (NSCs) generate new neurons that integrate into existing hippocampal networks and modulate mood and memory. These NSCs are largely quiescent and are stimulated by niche signals to activate and produce neurons. Wnt/β-catenin signalling acts at different steps along the hippocampal neurogenic lineage and has been shown to promote the proliferation of intermediate progenitor cells. However, whether it has a direct role in the regulation of NSCs still remains unclear. Here we used Wnt/β-catenin reporters and transcriptomic data from *in vivo* and *in vitro* models to show that both active and quiescent adult NSCs respond to Wnt/β-catenin signalling. Wnt/β-catenin stimulation instructed neuronal differentiation of active NSCs and promoted the activation or differentiation of quiescent NSCs in a dose-dependent manner. However, we found that inhibiting NSCs response to Wnt, by conditionally deleting β-catenin, did not affect their activation or maintenance of their stem cell characteristics. Together, our results indicate that whilst NSCs do respond to Wnt/β-catenin stimulation in a dose-dependent and state-specific manner, Wnt/β-catenin signalling is not cell-autonomously required to maintain NSC homeostasis, which could reconcile some of the contradictions in the literature as to the role of Wnt/β-catenin signalling in adult hippocampal NSCs.

## Introduction

Adult neurogenesis, the continuous process of generating new neurons from neural progenitors throughout life, is predominately restricted to two adult brain regions in mice: the ventricular-subventricular zone (V-SVZ) of the lateral ventricles and the dentate gyrus (DG) of the hippocampus, where adult born neurons contribute to olfactory bulb and hippocampal functions respectively (Bond et al., 2015; Lim et al., 2016; Deng et al., 2010). Adult hippocampal neurogenesis progresses via a distinct neurogenic lineage that begins with a population of multipotent radial glial-like neural stem cells (NSCs) that have both neurogenic and gliogenic potential (Steiner et al., 2004; Encinas et al., 2011; Bonaguidi et al., 2011; Pilz et al., 2018). The majority of NSCs remain out of the cell cycle in a reversible quiescent state (Doetsch et al., 1999; Andersen et al., 2014; Codega et al., 2014; Urbán et al., 2016). Adult neurogenesis is tightly controlled, with an important regulatory step being the transition of NSCs from quiescence to activation (reviewed by (Kempermann et al., 2015)). Too little stem cell activation results in an insufficient number of new neurons being generated (Andersen et al., 2014). On the other hand, excessive activation generates a transient burst in neurogenesis followed by a sharp decline resulting from the depletion of NSCs due to their limited self-renewal capacity (Paik et al., 2009; Renault et al., 2009). Tight regulation of the transition between quiescent and active NSC states is therefore crucial to ensure the long-term maintenance of the NSC pool while also meeting the neurogenic and gliogenic demands of the hippocampus. Niche-derived signals play an important role in regulating the transition between quiescent and active NSC states (Fuentealba et al., 2012; Choe et al., 2016; Imayoshi et al., 2010; Lie et al., 2005; Mira et al., 2010; Petrova et al., 2013; Qu et al., 2010). For example, Notch and BMP signalling have an integral role in regulating and maintaining NSC quiescence (Lavado et al., 2014; Ables et al., 2010; Bonaguidi et al., 2008; Mira et al., 2010). However, less is known about the niche derived signals that promote the activation of quiescent NSCs.

Wnt ligands and Wnt antagonists are expressed by multiple cell types within the DG niche, including the NSCs themselves (Lie et al., 2005; Qu et al., 2013; Jang et al., 2013; Seib et al., 2013). In the absence of a Wnt signal, β-catenin is phosphorylated for degradation by GSK3β, a component of the cytoplasmic β-catenin destruction complex (Daugherty et al., 2007). Upon Wnt ligand binding, activity of the destruction complex is impaired, allowing β-catenin stabilisation and translocation to the nucleus, where it forms a complex with TCF/LEF transcription factors to activate Wnt/β-catenin target gene expression, such as Axin2 (Lustig et al., 2002; Nusse et al., 2017). Wnt/β-catenin signalling is differentially active in cells along the neurogenic lineage and has been implicated in regulating both progenitor proliferation and newborn neuron maturation (Lie et al., 2005; Qu et al., 2013; Wexler et al., 2009; Kuwabara et al., 2009; Seib et al., 2013; Jang et al., 2013; Heppt et al., 2020; Rosenbloom et al., 2020).

Stimulating Wnt/β-catenin signalling in the DG promotes proliferation and neurogenesis (Lie et al., 2005; Seib et al., 2013; Jang et al., 2013). The absence of the Wnt inhibitor Dkk1 results in a specific increase in the generation of neuronally-committed intermediate progenitor cells (Seib et al., 2013). However, the absence of another Wnt inhibitor, sFRP3, stimulates proliferation but does not favour a neurogenic lineage choice of NSCs (Jang et al., 2013). Inhibiting Wnt/β-catenin signalling has also yielded contrasting results, as it has been shown to both impair the generation of newborn neurons *in vivo* and induce neuronal differentiation *in vitro* (Qu et al., 2013; Qu et al., 2010; Wexler et al., 2009; Kuwabara et al., 2009; Lie et al., 2005). These discrepancies could be due to the use of different experimental approaches to modulate Wnt/β-catenin signalling, such as overexpressing a Wnt ligand vs deleting a Wnt inhibitor, or to the use of *in vitro* vs *in vivo* approaches (Lie et al., 2005; Seib et al., 2013; Jang et al., 2013; Kuwabara et al., 2009; Wexler et al., 2009). The majority of these studies have used systemic or hippocampal-wide modulation of Wnt/β-catenin signalling, which do not discriminate between cell autonomous and non-autonomous effects, neither does it allow identification of the step(s) in the NSC lineage when Wnt/β-catenin signalling acts (Lie et al., 2005; Seib et al., 2013; Jang et al., 2013; Qu et al., 2013). Also, the use of constitutive knock-out mice makes it difficult to distinguish between a developmental and an adult neurogenesis phenotype (Seib et al., 2013; Jang et al., 2013; Qu et al., 2013; Qu et al., 2010). Furthermore, earlier studies did not robustly distinguish between NSCs with radial morphology (NSCs) and intermediate progenitor cells (IPCs), and therefore the role of Wnt/β-catenin signalling in NSCs, and particularly in their transition between active and quiescent states is still unclear (Kuwabara et al., 2009; Seib et al., 2013).

Here we have used genetic and pharmacological tools to measure and manipulate Wnt/β-catenin signalling specifically in active and quiescent adult NSCs, both *in vivo* and *in vitro*. We show that both quiescent and active NSCs respond to Wnt/β-catenin signalling; that the response of NSCs to Wnt/β-catenin stimulation is dose- and cell-state-specific; but that Wnt/β-catenin signalling is not essential for cell-autonomous NSC homeostasis or for their ability to proliferate and generate early neuronal progenitors. Together, these findings reconcile some of the current contradictions as to the role of Wnt/β-catenin signalling in adult hippocampal NSCs.

## Results

### Quiescent and active NSCs show similar levels of Wnt/β-catenin signalling activity *in vivo*

To study the role of Wnt/β-catenin in the transition of hippocampal stem cells from quiescence to activation, we first investigated whether quiescent and active hippocampal stem cells express components of the Wnt/β-catenin signalling pathway that would allow them to transduce a Wnt signal. Expresssion of Wnt/β-catenin signalling components in adult hippocampal NSCs has been reported in previously published single cell sequencing datasets, but in most too few NSCs were identified to allow a robust comparison between quiescent and active states (Hochgerner et al., 2018). We re-analysed a previously generated single-cell RNA sequencing dataset containing 2,947 NSCs (Fig. 1A) (Harris et al., 2020). We found that both quiescent and active NSCs heterogeneously express components of the Wnt/β-catenin signalling pathway (Fig. 1A). Both quiescent and active NSCs express Wnt ligands *(Wnt7a* and *Wnt7b)* as well as Wnt inhibitors *(Dkk3* and *sFRP1*), suggesting that NSCs directly regulate Wnt activity levels in the DG (Fig. 1A).

**Figure 1:**
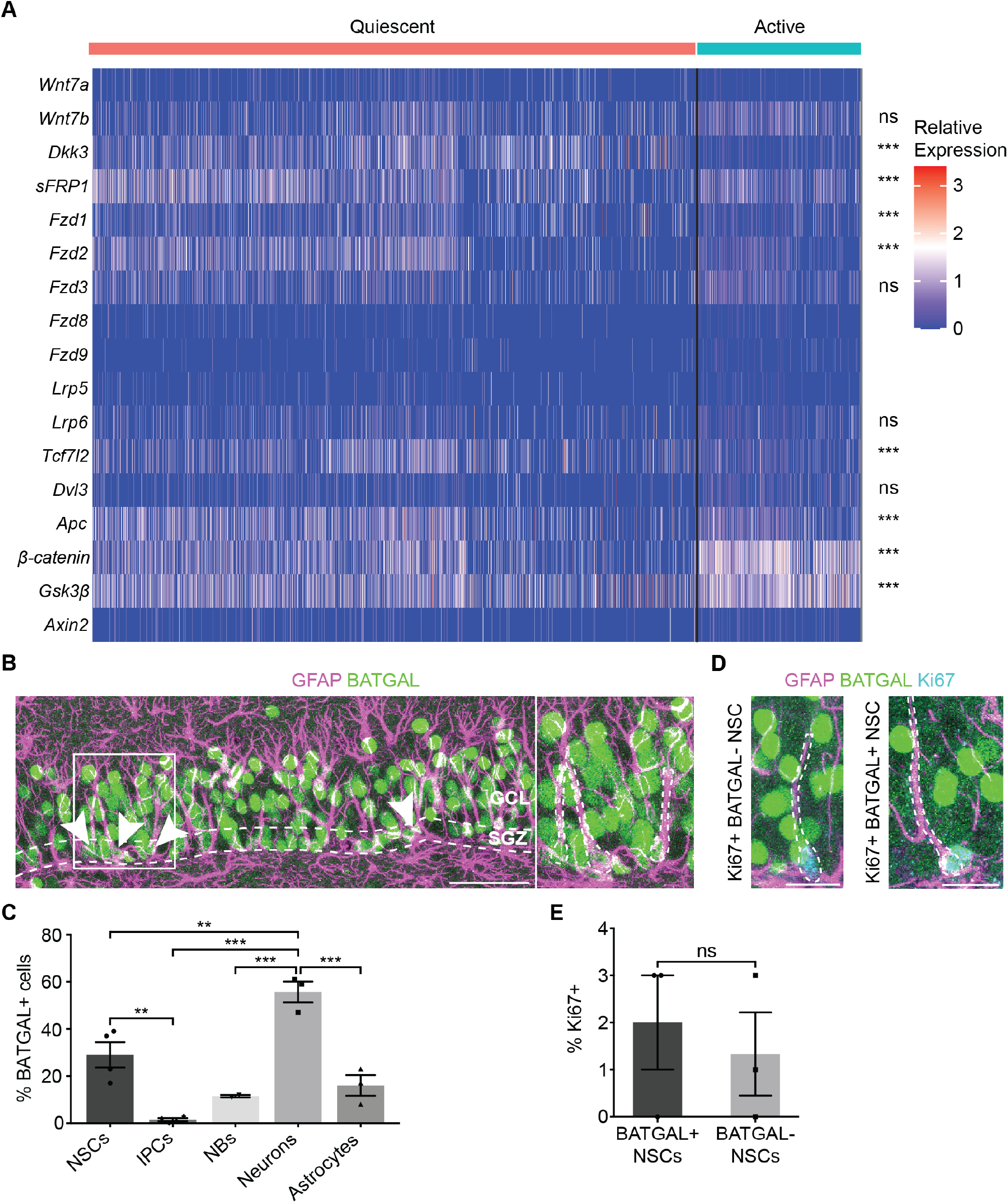
NSCs *in vivo* respond to Wnt/β-catenin signalling independently of their activation state. (**A**) Heatmap representation of publicly available (GSE159768) single cell RNA sequencing data showing the expression of Wnt/β-catenin pathway components from *in vivo* hippocampal quiescent and active NSCs as previously described (Harris et al., 2020). (**B**) BATGAL (β-galactosidase) and GFAP immunolabelling in the DG of 2-month-old BATGAL Wnt/β-catenin reporter mouse (Maretto et al., 2003). Arrowheads indicate BATGAL+ NSCs (SGZ cell body, GFAP+ radial process). Region in white box is enlarged in adjacent panel to show three BATGAL+ NSCs. Dashed lines denote the SGZ. SGZ, Subgranular zone. GCL, Granule cell layer. Scale bar, 50μm. (**C**) Quantification of the proportion of BATGAL+ cells in the DG of 2-month-old BATGAL mice. NSCs (GFAP+ NSCs, 29 ± 5.44%, n=4), IPCs (GFAP-Sox2+, 1.5 ± 0.6%, n=4), neuroblasts (NBs, DCX+, 11.5 ± 0.5, n=2), neurons (NeuN+, 55.67 ± 4.44%, n=3), astrocytes (GFAP+ Sox2-, 16 ± 4.44%, n=3). (**D**) BATGAL (β-galactosidase) immunolabelling in Ki67+ NSCs in 2-month-old BATGAL mice. Ki67+ BATGAL+ NSCs and Ki67+ BATGAL-NSCs are outlined. Scale bar, 20μm. (**E**) Quantification of the proportion of Ki67+ immunolabelling in BATGAL+ (2 ± 1%) and BATGAL-NSCs (1.33 ± 0.99%) in (**D**). Proliferation is not enriched in BATGAL+ NSCs. n=3. Statistics: Student’s t-test (**A**), ordinary one-way ANOVA with Tukey’s multiple comparisons test (**C**) and an unpaired Student’s t-test (**E**). (ns, p>0.05. **, p<0.01. ***, p<0.001). Error bars represent mean with SEM.

Quiescent NSCs express Wnt receptors *(Fzd1* and *Fzd2)* at a higher level compared with active NSCs, which corroborates previous reports showing that quiescent NSCs are enriched for cell surface receptors (Fig. 1A) (Shin et al., 2015; Hochgerner et al., 2018; Artegiani et al., 2017; Cheung et al., 2013). The Wnt transducer molecule *Tcf7l2* is also higher in quiescent NSCs than in active ones, while *β-catenin* and *GSK3β* are both upregulated in active NSCs (Fig. 1A). These differences in expression of Wnt related genes, however, do not translate into a differential expression of Wnt targets between quiescent and active NSCs. *Axin2* is expressed at low levels by very few quiescent (5%) and active (7%) NSCs (Fig. 1A), which could suggest that few NSCs respond to Wnt/β-catenin signalling at any given time, but could also reflect the limitation of single cell RNA sequencing in detecting lowly expressed genes (Haque et al., 2017). Nevertheless, the similar *Axin2* expression levels between quiescent and active NSCs, suggests that cells in these two states respond similarly to Wnt/β-catenin signalling (Fig. 1A).

To further characterise the response of NSCs and other DG cells to Wnt/β-catenin signalling *in vivo,* we next used the BATGAL Wnt/β-catenin reporter mouse model, which expresses *β-galactosidase* under the control of 7xTCF/LEF binding sites (Maretto et al., 2003). We identified NSCs based on their glial fibrillary acidic protein (GFAP) expression, their subgranular zone (SGZ) cell body localisation and their radial process extending through the granule cell layer (GCL) of the DG (Fig. 1B). By co-immunolabelling the DG of two-month-old BATGAL mice for β-galactosidase alongside cell-specific markers, we found that a higher proportion of NSCs (29±5.4%) and neurons (55.7±4.4%) respond to Wnt/β-catenin signalling compared with IPCs (1.5±0.6%), neuroblasts (11.5±0.5%) and astrocytes (16±4.4%) (Fig. 1C). This corroborates previous reports and validates the use of BATGAL mice to characterise the Wnt/β-catenin response in NSCs (Heppt et al., 2020; Garbe et al., 2012). To investigate whether the Wnt/β-catenin response correlates with NSC activation states, we quantified proliferation in BATGAL-positive and -negative NSCs (Fig. 1D). The proportion of proliferating NSCs was similar between BATGAL-positive or BATGAL-negative NSCs. (Fig. 1E). Overall, these data show that NSCs *in vivo* respond similarly to Wnt/β-catenin signalling, independently of their activation state.

### NSC maintenance and adult neurogenesis are unaffected by deletion of β-catenin in NSCs *in vivo*

To investigate the effects of Wnt/β-catenin inhibition in NSCs *in vivo*, we generated *GlastCreERT2; β-cat^fl/fl ex3-6^; RYFP* mice to conditionally delete β-catenin in Glast-expressing NSCs by tamoxifen-inducible, Cre-mediated recombination (Huelsken et al., 2001; Mori et al., 2006; Srinivas et al., 2001). However, we observed that the β-catenin allele failed to recombine in NSCs (Fig. S1A-C), therefore we generated a second β-catenin floxed mouse line; *GlastCreERT2; β-cat^fl/fl ex2-6^; RYFP* mice (hereafter referred to as β-cat^del ex2-6^ mice) that contained a different conditional mutant allele of β-catenin *(β-cat^fl/fl ex2-6^* allele, Fig. S1D) (Brault et al., 2001). The expression of the recombined *β-catenin* transcript was significantly decreased in YFP-positive FAC-sorted cells from the DG of β-cat^del ex2-6^ mice compared with control (Fig. S1E, F), indicating successful recombination of the *β-cat^fl/fl-ex2-6^* allele. We crossed β-cat^del ex2-6^ mice with BATGAL mice to examine the Wnt/β-catenin response in recombined NSCs and found that the proportion of BATGAL-positive NSCs was reduced in β-cat^del ex2-6^ mice compared with control 30 days after tamoxifen administration in two-month-old-mice, confirming that loss of β-catenin impairs Wnt/β-catenin activity in NSCs (Fig. S1G-I). To examine the effect of β-catenin deletion on NSCs, we administered tamoxifen to two-month-old β-cat^del ex2-6^ and control mice and performed immunofluorescence analysis 30 days later (Fig. 2A-B). The proportion of Ki67-positive NSCs was not significantly different between β-cat^del ex2-6^ and control mice, indicating that β-catenin deletion did not perturb NSC proliferation (Fig. 2C). We also quantified the total number of NSCs and found no significant difference between β-cat^del ex2-6^ and control mice (Fig. 2D), indicating that NSC maintenance was unaffected by β-catenin deletion. Overall, these data suggest that the proliferation and maintenance of NSCs are unaffected by loss of Wnt/β-catenin signalling.

**Figure 2:**
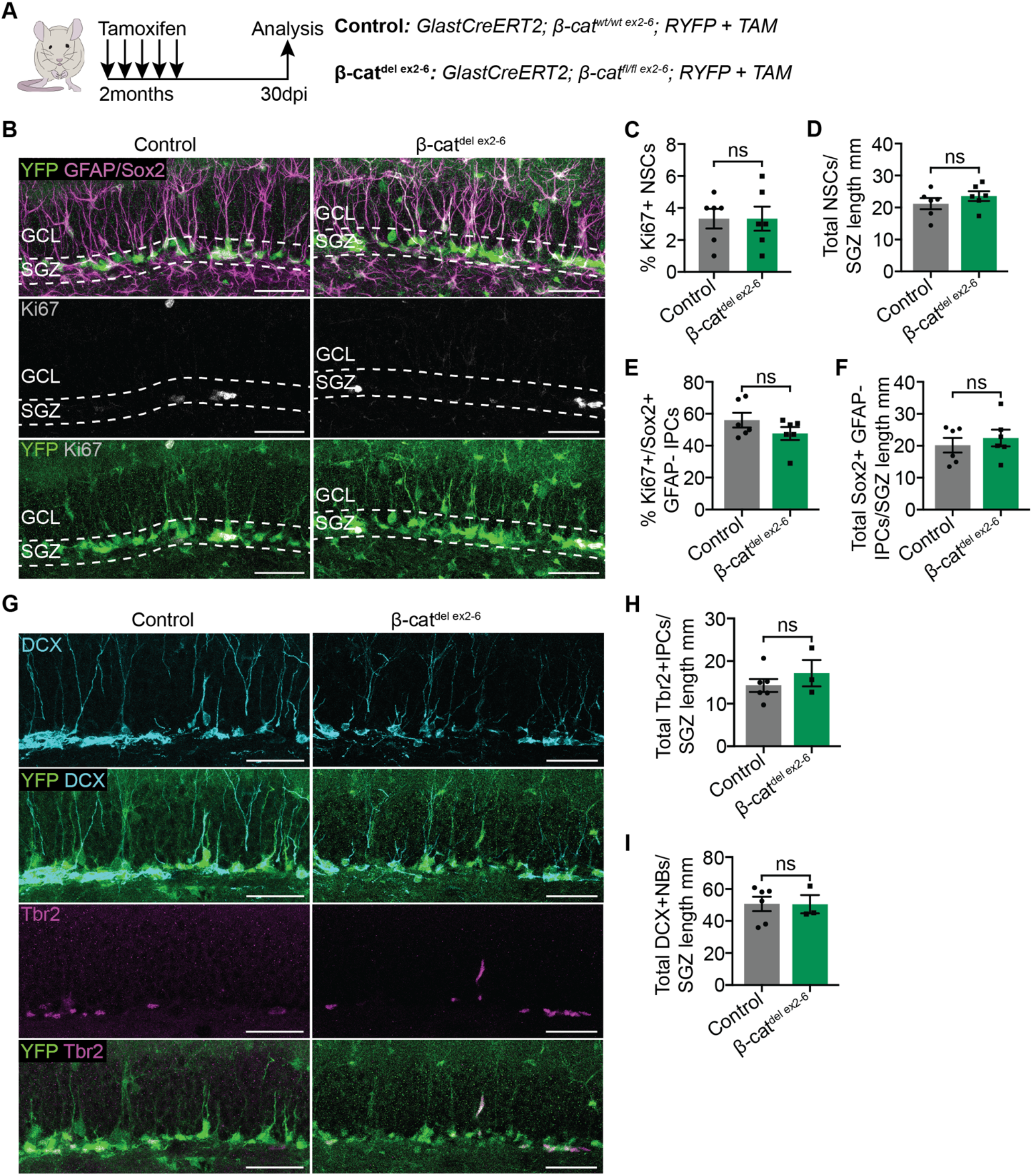
NSCs and adult hippocampal neurogenesis are unaffected by the NSC specific deletion of β-catenin and inhibition of Wnt/β-catenin signalling. (**A**) Two-month-old Control and β-cat^del ex2-6^ mice were administered tamoxifen for 5 consecutive days and sacrificed 30 days after the first tamoxifen injection. (**B**) YFP, GFAP, Sox2 and Ki67 immunolabelling in the DG of Control and β-cat^del ex2-6^ mice 30 days after tamoxifen administration. Scale bars, 50μm. (**C**-**F**) Quantifications of the data shown in (**B**). The proportion of Ki67+ NSCs (Control vs β-cat^del ex2-6^: 3.33 ± 0.61% vs 3.333 ± 0.76%, **C**), the total number of NSCs (YFP+ GFAP+ Sox2+ radial cells in the SGZ) normalised to the length of the SGZ (mm) (Control vs β-cat^del ex2-6^: 21.2 ± 1.73 vs 23.56 ± 1.52, **D**), the proportion of proliferating (Ki67+) IPCs (Sox2+ YFP+ GFAP-cells in the SGZ; Control vs β-cat^del ex2-6^: 56 ± 4.56% vs 47.67 ± 4.03%, **E**) and the total number of IPCs normalised to the length of the SGZ (mm) (Sox2+ YFP+ GFAP-cells in the SGZ; Control vs β-cat^del ex2-6^: 20.18 ± 2.31 vs 22.43 ± 2.62, **F**) are unchanged between Control and β-cat^del ex2-6^ mice, indicating that NSCs and IPCs are unaffected by β-catenin deletion. n=6. (**G**) YFP, Tbr2 and DCX immunolabelling in the DG of Control and β-cat^del ex2-6^ mice 30 days after tamoxifen administration. Scale bars, 50μm. (**H, I**) Quantifications of the data shown in (**G**). The total number of Tbr2+ IPCs normalised to the SGZ length (mm) (Tbr2+ YFP+ cells in the SGZ; Control vs β-cat^del ex2-6^: 14.29 ± 1.51 vs 17.16 ± 3.09, **H**) and the total number of neuroblasts (NBs) normalised to the SGZ length (mm) (DCX+ YFP+ cells; Control vs β-cat^del ex2-6^: 50.72 ± 4.49 vs 50.53 ± 5.69, **I**) are unchanged between Control and β-cat^del ex2-6^ mice, indicating that Tbr2+ IPCs and NBs are unaffected by β-catenin deletion. n=6 and n=3 for Control and β-cat^del ex2-6^ mice respectively. Statistics: unpaired Student’s t-test (**C-F, H-I**). (ns, p>0.05). Error bars represent mean with SEM.

We then investigated how the loss of β-catenin affects later steps in the adult hippocampal neurogenic lineage. We quantified the percentage of proliferating IPCs (Ki67-positive, Sox2-positive, GFAP-negative cells) (Fig. 2E), the total number of IPCs, identified either as Sox2-positive/GFAP-negative cells (Fig. 2F) or Tbr2-positive cells (Fig. 2G, H), and the total number of neuroblasts, identified by doublecortin (DCX) staining (Fig. 2G, I), and did not find any significant difference between β-cat^del ex2-6^ and control mice. Overall, these data show that loss of Wnt/β-catenin signalling in NSCs *in vivo* does not impair the behaviour or maintenance of NSCs nor does it impair the generation of neuronally-committed precursors in the adult hippocampus.

### Stabilising β-catenin in NSCs causes their displacement and loss from the DG niche

After having assessed the effects of disrupting Wnt/β-catenin signalling in NSCs *in vivo*, we wanted to investigate how stimulating Wnt/β-catenin signalling would affect NSCs. For this, we generated an *in vivo* mouse model, *GlastCreERT2; Catnb^lox(ex3)/wt^ RYFP* mice (hereafter referred to as Catnb^del(ex3)^ mice), to conditionally stabilise β-catenin in Glast-expressing NSCs following tamoxifen-inducible Cre-mediated recombination of the *Catnb^lox(ex3)^* allele (Harada et al., 1999). The *Catnb^lox(ex3)^* allele is a conditional constitutively active allele of *β-catenin* where exon 3, which encodes the GSK3β phosphorylation sites that mark β-catenin for degradation, is flanked by LoxP sites (Harada et al., 1999). Therefore, Cre-mediated recombination of the *Catnb^lox(ex3)^* allele results in β-catenin stabilisation and ligand-independent activation of downstream Wnt/β-catenin signalling in targeted cells. Upon recombination, we observed an increase in β-catenin levels as well as an increase in the intensity of the BATGAL reporter in Catnb^lox(ex3)^/BATGAL mice compared to BATGAL controls (Fig. S2B-E).

We injected 2-month-old Catnb^del(ex3)^ and control mice with tamoxifen for 5 consecutive days and analysed their NSCs 10 and 30 days later (Fig. S2A). We first assessed the effect of stabilising β-catenin in NSCs by quantifying the total number of NSCs in Catnb^del(ex3)^ and control mice (Fig. S2 F-K). We found a significant decrease in number of NSCs in Catnb^del(ex3)^ mice compared with control at 30 days (Fig. S2G) but not at 10 days after tamoxifen administration (Fig. S2J), suggesting that stabilising β-catenin causes NSC loss between 10 and 30 days later. This could be due to increased NSC proliferation and subsequent depletion. Although 30 days after tamoxifen administration we observed an increase in the proportion of Ki67-positive NSCs (Fig. S2H), we did not see this at 10 days after tamoxifen administration (Fig. S2K). Therefore, β-catenin stabilisation does not cause an immediate increase in proliferation. The loss of NSCs at 30 days could still be due to increased proliferation between the 10 and 30 day time points, however we noticed that the cellular organisation of the DG was disrupted in Catnb^del(ex3)^ mice already at 10 days after β-catenin stabilisation. Many YFP-positive recombined cells were displaced throughout the GCL and molecular layer (ML) (Fig. S2I). Some of the displaced cells retained NSC characteristics (GFAP-positive, Sox2-positive cells with radial morphology) but their cell bodies were not correctly located in the SGZ (Fig. S2L). Such cells were also present in control mice, but the proportion of displaced NSCs was increased threefold in Catnb^del(ex3)^ compared with control mice although this was highly variable between Catnb^del(ex3)^ mice (Fig. S2M, N). This suggests that stabilising β-catenin in NSCs promotes their displacement from their correct SGZ location, which might cause their subsequent loss from the DG. As β-catenin also regulates cell adhesion at adherens junctions (Bienz. 2005), this displacement phenotype could be due to disrupted cell adhesion, which precludes using this mouse model to investigate the effects of stimulating Wnt/β-catenin signalling in NSCs *in vivo*. As an alternative approach, we next decided to use an established *in vitro* model of hippocampal NSCs that allows manipulation of their quiescent and active states in a niche-independent setting (Blomfield et al., 2019).

### Quiescent and active NSCs *in vitro* show similar levels of Wnt/β-catenin signalling activity

Dissociated hippocampal NSCs in adherent cultures can be maintained in a proliferative state by the presence of FGF2 and can be induced into a reversible state of quiescence through the addition of BMP4 (Blomfield et al., 2019; Martynoga et al., 2013; Mira et al., 2010). Using a published bulk RNA sequencing dataset (Blomfield et al., 2019), we found that NSCs in this *in vitro* model system largely recapitulate the expression of Wnt receptors and pathway components observed in quiescent and active NSCs in *vivo* (Fig. 3A), like for instance the upregulation of Wnt receptors in quiescent NSCs compared with active NSCs, which also corroborates published reports (Lie et al., 2005; Wexler et al., 2009). Moreover, quiescent and active NSCs express Wnt ligands at both the RNA and protein levels (Fig. 3A, B), suggesting they can self-regulate their behaviour by autocrine/paracrine Wnt/β-catenin signalling as previously proposed (Qu et al., 2013; Qu et al., 2010; Wexler et al., 2009). As with NSCs *in vivo,* we found that quiescent and active NSCs *in vitro* express *Axin2* at similar levels, thus suggesting that their response to Wnt/β-catenin signalling is independent of their activation state (Fig. 3A).

**Figure 3:**
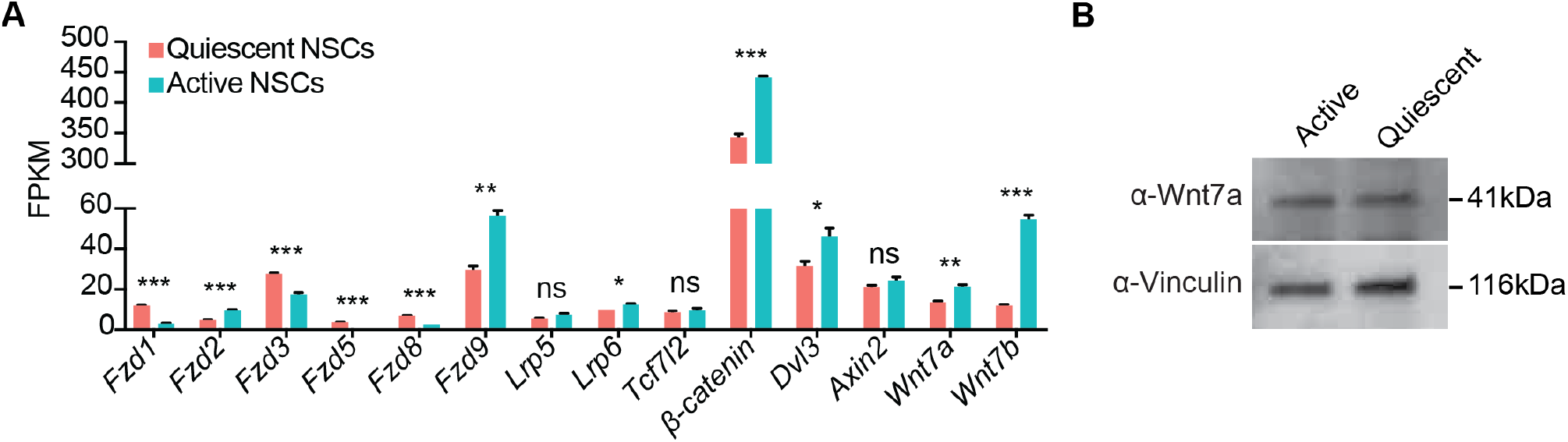
NSCs in quiescent and active culture conditions express components of the Wnt/β-catenin signalling pathway. (**A**) Gene expression data from publicly available (GSE116997) bulk RNA sequencing analysis comparing the expression levels of Wnt receptors *(Fzd1, Fzd2, Fzd3, Fzd5, Fzd8, Fzd9, Lrp5* and *Lrp6*), transducer molecules *(Tcf7l2, β-catenin* and *Dvl*), the Wnt/β-catenin target gene *Axin2* and Wnt ligands *(Wnt7a* and *Wnt7b)* in quiescent and active NSC cultures. n=3. (**B**) Wnt7a Western blot in active and quiescent NSC lysates shows that active and quiescent NSCs produce Wnt protein. Statistics: unpaired Student’s t-test (**A**). (ns, p>0.05. *, p<0.05. **, p<0.01. ***, p<0.001). Error bars represent mean with SEM.

### Loss of Wnt/β-catenin signalling has no effect on the maintenance of NSC activation states or stem cell potency *in vitro*

We next took advantage of this *in vitro* system to investigate the role of Wnt/β-catenin signalling specifically in active or quiescent NSCs as well as in their transition between states and to determine whether Wnt/β-catenin is dispensable for NSCs *in vitro*, as we demonstrated *in vivo*. We derived an NSC cell line from *β-cat^fl/fl ex3-6^*; RYFP mice (Huelsken et al., 2001; Srinivas et al., 2001), hereafter referred to as β-cat^del ex3-6^ NSCs. When β-cat^del ex3-6^ NSCs were transduced with a Cre-expressing adenovirus, Cre-recombined cells were identified by their expression of YFP. To confirm successful recombination of the *β cat^fl/fl ex3-6^* allele (Fig. S1A) at various time points after Cre-transduction, we performed qPCR and Western blot analysis (Fig. 4A and Fig. S3A, B). Expression of the recombined portion of the *β*-cat transcript was significantly decreased by 24hrs after Cre-transduction and β-catenin protein became undetectable by 48hrs, indicating successful recombination of the *βcat^fl/fl ex3-6^* allele (Fig. S3A, B). We next examined *Axin2* expression levels in β-cat^del ex3-6^ and control NSCs treated with the Wnt/β-catenin agonist CHIR99021, which stimulates Wnt/β-catenin signalling by inhibiting GSK3β of the destruction complex (Ring et al., 2003). CHIR99021 treatment upregulated *Axin2* expression levels in control NSCs but not in β-cat^del ex3-6^ NSCs, confirming that deletion of β-catenin impairs the response of NSCs to Wnt/β-catenin signalling (Fig. S3C).

**Figure 4:**
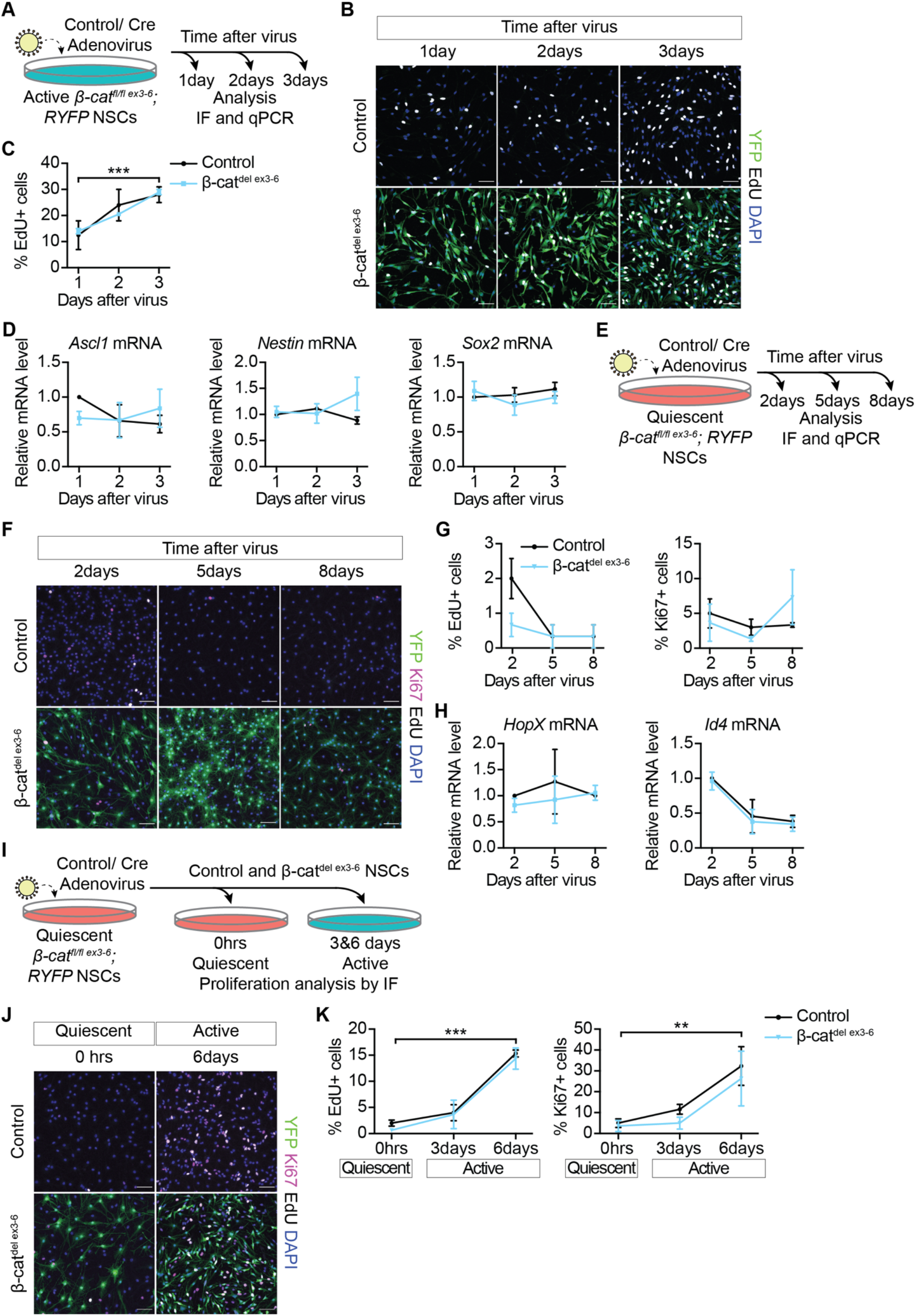
NSC behaviours are unaffected by loss of β-catenin and impaired Wnt/β-catenin signalling *in vitro.* (**A**) Active β-cat^fl/fl ex3-6^ NSCs were transduced with Control or Cre-adenovirus and samples collected 1-, 2- and 3-days later. (**B**) YFP, β-catenin, EdU and DAPI immunolabelling 1-, 2- and 3-days after Control- and Cre-adenovirus transduction in Control and β-cat^del ex3-6^ active NSCs. YFP immunolabelling identifies recombined β-cat^del ex3-6^ NSCs. Scale bars, 50μm. (**C**) Quantification of the proportion of EdU+ cells in (**B**). Proliferation is unchanged in Control and β-cat^del ex3-6^ active NSCs (Control vs β-cat^del ex3-6^: 1day = 12.5 ± 5.5% vs 14 ± 1%, 2days = 24 ± 6% vs 20.5 ± 0.5%, 3days = 28 ± 3% vs 29 ± 1%). In β-cat^del ex3-6^ active NSCs the proportion of EdU+ cells was calculated as a fraction of the YFP+ recombined cells. The increase in the proportion of proliferating cells is statistically significant across time. n=2. (**D**) Expression of the stem cell associated genes *Ascl1, Nestin* and *Sox2* are unchanged in β-cat^del ex3-6^ active NSCs compared with Control 1-, 2- and 3-days after virus transduction. n=3. (**E**) Quiescent β-cat^fl/fl ex3-6^ NSCs were transduced with control or Cre-adenovirus following 72hrs quiescence induction and samples collected 2-, 5- and 6-days later. (**F**) YFP, EdU, Ki67 and DAPI immunolabelling 2-, 5- and 8-days after control- and Cre-adenovirus transduction in Control and β-cat^del ex3-6^ quiescent NSCs. Scale bars, 50μm. (**G**) Quantification of the proportion of EdU+ cells (Control vs β-cat^del ex3-6^: 2days = 2 ± 0.58% vs 0.67 ± 0.33%, 5days = 0.33 ± 0.33% vs 0.33 ± 0.33%, 8days = 0.33 ± 0.33% vs 0.33 ± 0.33%) and Ki67+ cells in (**D**) (Control vs β-cat^del ex3-6^: 2days = 5 ± 2.08% vs 3.67 ± 2.67%, 5days = 3 ± 1.16% vs 1.33 ± 0.33%, 8days = 3.33 ± 0.33% vs 7.33 ± 3.93%). Proliferation of quiescent NSCs is unchanged by the loss of β-catenin. n=3. (**H**) Expression of the quiescence associated genes *HopX* and *Id4* is unchanged by the loss of β-catenin. n=3. (**I**) Two-days after virus transduction of quiescent Control and β-cat^fl/fl ex3-6^ NSCs, samples were collected before (0hrs) and at 3- and 6-days after retuning cells to proliferative culture conditions for immunofluorescence analysis of proliferation markers. (**J**) YFP, EdU, Ki67 and DAPI immunolabelling in quiescent (0hrs) and 3- and 6-days after returning Control and β-cat^del ex3-6^ NSCs to proliferative culture conditions. Scale bars, 50μm. (**K**) Quantification of the data shown in (**J**) of the proportion of EdU+ cells (Control vs β-cat^del ex3-6^: 0hrs = 2 ± 0.58% vs 0.67 ± 0.33%, 3days = 4 ± 1.53% vs 3.67 ± 2.73%, 6days = 15.33 ± 0.67% vs 14.33 ± 2.03%) and Ki67+ cells (Control vs β-cat^del ex3-6^: 0hrs = 5 ± 2.08% vs 3.67 ± 2.67%, 3days = 11.67 ± 2.33% vs 5 ± 2.89%, 6days = 32.33 ± 9.28% vs 26.33 ± 13.09%). Both Control and β-cat^del^ ^ex3-6^ quiescent NSCs reactivate similarly when returned to proliferative culture conditions. The increase in the proportion of proliferating cells is statistically significant across time. n=3. Statistics: Two-way ANOVA with Sidak’s multiple comparisons test (**C, D, G, H, K**). (ns, p>0.05. *, p<0.05, **, p<0.01. ***, p<0.001). Error bars represent mean with SEM.

We then asked whether the loss of β-catenin, and the resulting inhibition of Wnt/β-catenin signalling, affects the stem cell identity and proliferation of NSCs. To investigate proliferation changes, control and β-cat^del ex3-6^ active NSCs were pulsed with EdU for 1 hour to label cells in S-phase of the cell cycle at different time points after Cre-transduction (Fig. 4A). The proportion of EdU-positive cells was unchanged between control and β-cat^del ex3-6^ active NSCs at all time points analysed, indicating that active NSC proliferation is unaffected by loss of β-catenin (Fig. 4B, C). The stem cell-associated genes, *Ascl1, Nestin* and *Sox2* were similarly expressed between β-cat^del ex3-6^ and control active NSCs at all time points (Fig. 4D), suggesting that loss of β-catenin does not affect the maintenance of NSC identity under these conditions.

We next assessed how the loss of Wnt/β-catenin signalling affects the ability of NSCs to maintain quiescence. To do this, we repeated the previous experiment but cultured the β-cat^del ex3-6^ and control NSCs under quiescent conditions (Fig. 4E). We found that both cultures maintained a similarly low rate of proliferation over time (Fig. 4F, G). This suggests that maintenance of quiescence is unaffected by the loss of Wnt/β-catenin signalling. Indeed, the quiescence-associated genes *HopX* and *Id4* were similarly expressed in control and β-cat^del ex3-6^ quiescent NSCs (Fig. 4H). Overall, these data suggest that BMP-induced NSC quiescence is unaffected by inhibition of Wnt/β-catenin signalling.

So far, we have shown that Wnt/β-catenin signalling is not required to maintain established active or quiescent states of NSCs. However, the role of Wnt/β-catenin signalling in enabling the transition between states, i.e. the activation of quiescent NSCs, remains unaddressed. To investigate this, we returned β-cat^del ex3-6^ and control quiescent NSCs to proliferative culture conditions to promote reactivation (Fig. 4I). We measured proliferation immediately before (time zero) and at three and six days after returning the cells to proliferative culture conditions (Fig. 4I, J). Proliferation increased similarly in both β-cat^del ex3-6^ and control NSCs, as shown by comparable increases in the proportions of EdU-positive and Ki67-positive cells (Fig. 4K). This suggests that loss of intact Wnt/β-catenin signalling does not affect the transition of NSCs from quiescent to active states.

As NSC proliferation and activity are not affected by loss of Wnt/β-catenin signalling, we next tested whether neuronal or astrocytic differentiation was affected by the chronic loss of β-catenin. We changed the media conditions of β-cat^del ex3-6^ and control active NSCs at 6-, 12- and 18-days after Cre-transduction to promote neuronal differentiation (B27 for 5 days) or astrocytic differentiation (FBS for 5 days) (Fig. S4A). We then immunolabelled cells for the neuronal marker MAP2 and astrocyte marker GFAP to identify and quantify the proportions of neurons and astrocytes, respectively (Fig. S4B-E). We found that β-cat^del ex3-6^ NSCs generated a similar proportion of MAP2-positive neurons to control NSCs (Fig. S4C). The proportions of GFAP-positive astrocytes generated were also similar between β-cat^del ex3-6^ and control NSCs (Fig. S4E). This suggests that the neurogenic and gliogenic potential of NSCs is unaffected by the loss of β-catenin dependent Wnt signalling under these differentiating conditions *in vitro*.

### Wnt/β-catenin stimulation promotes the neuronal differentiation of active NSCs

We next moved on to investigate the consequences of stimulating Wnt/β-catenin signalling in the NSC cultures. For this, we treated active NSCs with the GSK3β inhibitor CHIR99021 for 48hrs. CHIR99021 treatment resulted in a dose-dependent increase in *Axin2* expression levels, which were significantly higher compared with control cells with 5μM and 10μM CHIR99021 treatments (Fig. 5A).

**Figure 5:**
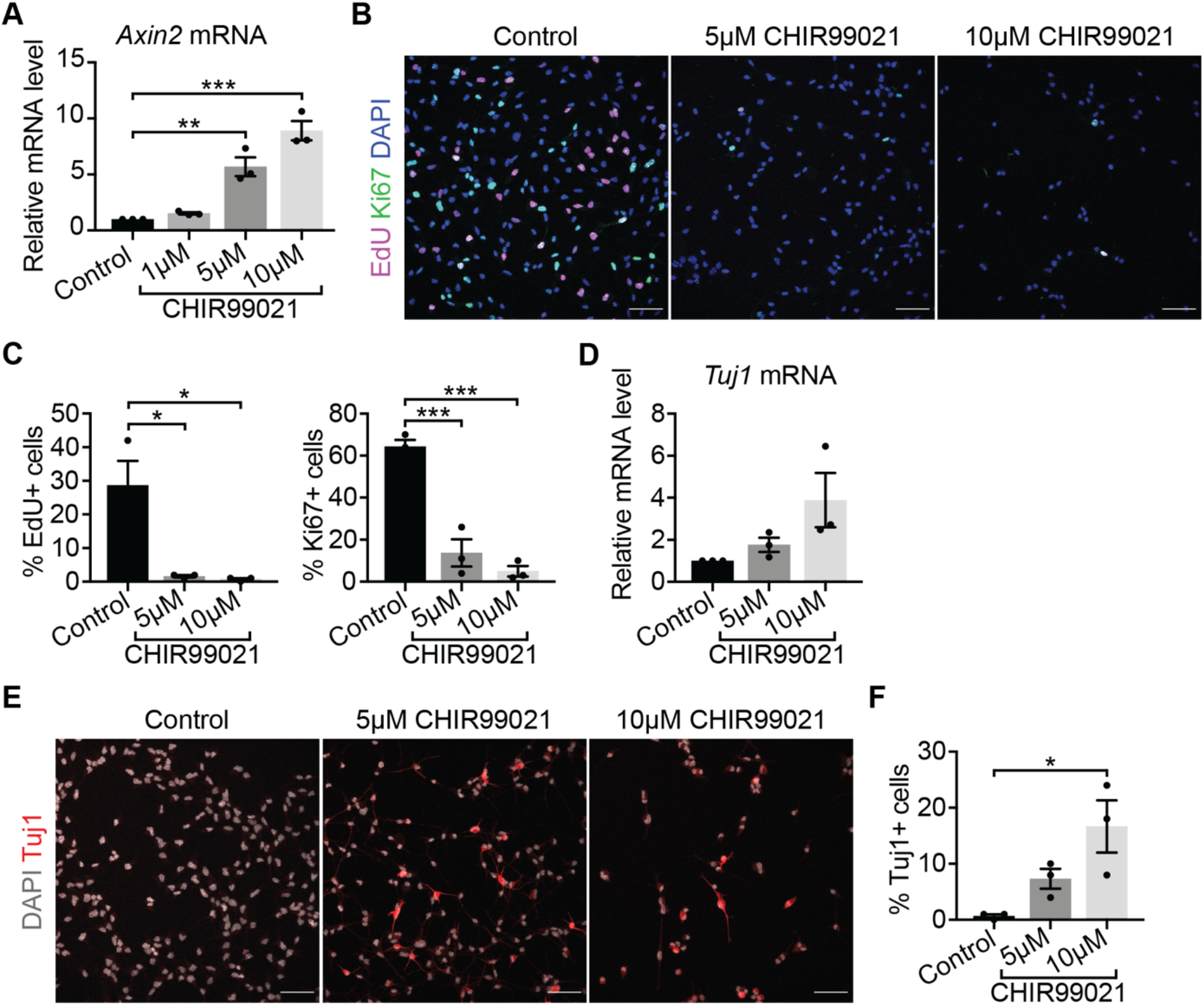
Stimulating Wnt/β-catenin signalling promotes neuronal differentiation of active NSCs. (**A**) Upregulation of *Axin2* expression in active NSCs treated with 5μM and 10μM CHIR99021 for 48hrs confirms stimulation of Wnt/β-catenin signalling. n=3. (**B**) Immunolabelling of proliferation markers EdU and Ki67 and DAPI in active NSCs treated with CHIR99021. Scale bars, 50μm. (**C**) Quantifications of the data shown in (**B**). The proportion of EdU+ cells (Control, 28.67 ± 7.27%. 5μM CHIR99021, 1.67 ± 0.33%. 10μM CHIR99021, 0.67 ± 0.33%) and of Ki67+ cells (Control, 64.33 ± 3.18%. 5μM CHIR99021, 13.67 ± 6.49%. 10μM CHIR99021, 5 ± 2.52%) is decreased upon CHIR99021 treatment. n=3. (**D**) *β-III-Tubulin (Tuj1)* expression is upregulated by 48hrs of CHIR99021 treatment in active NSCs. (10μM CHIR99021, p=0.06). n=3. (**E**) Immunolabelling of Tuj1 and DAPI in active NSCs treated with CHIR99021 compared to Control. Scale bars, 50μm. (**F**) Quantifications of the data shown in (**E**) of the proportion of Tuj1+ cells (Control, 0.67 ± 0.33%. 5μM CHIR99021, 7.33 ± 1.76%. 10μM CHIR99021, 16.67 ± 4.67%). n=3. Statistics: Repeated measures one-way ANOVA with Dunnett’s multiple comparison test (**A, C, D and F**). (ns, p>0.05. *, p<0.05. **, p<0.01. ***, p<0.001). Error bars represent mean with SEM.

We then investigated whether 5μM and 10μM CHIR99021 treatments led to changes in proliferation of active NSCs, and found that the proportions of EdU-positive and of Ki67-positive NSCs were significantly decreased in active NSCs treated with CHIR99021 compared with control (Fig. 5B, C). As reduced proliferation could be the result of an increased differentiation of CHIR99021-treated NSCs, we next examined the expression of the neuronal marker *β-III-tubulin (Tuj1;* Fig. 5E). We found that the expression of *Tuj1* as well as the proportion of Tuj1-positive immunolabelled cells was increased in active NSCs upon 10μM CHIR99021 treatment (Fig. 5D-F). Overall, these results show that stimulating Wnt/β-catenin signalling in active NSCs reduces their proliferative capacity and promotes neuronal differentiation.

### Wnt/β-catenin stimulation promotes the activation and differentiation of quiescent NSCs in a dose-dependent manner

To investigate the effect of stimulating Wnt/β-catenin signalling in quiescent NSCs, we similarly treated quiescent NSCs with 1μM CHIR99021, 5μM CHIR99021 and 10μM CHIR99021 for 48hrs. Using *Axin2* expression levels as a read-out of Wnt/β-catenin signalling levels, we found that, similarly to active NSCs, *Axin2* expression was upregulated in a dose-dependent manner by CHIR99021 in quiescent NSCs, starting at 5μM CHIR99021 treatment (Fig. 6A).

**Figure 6:**
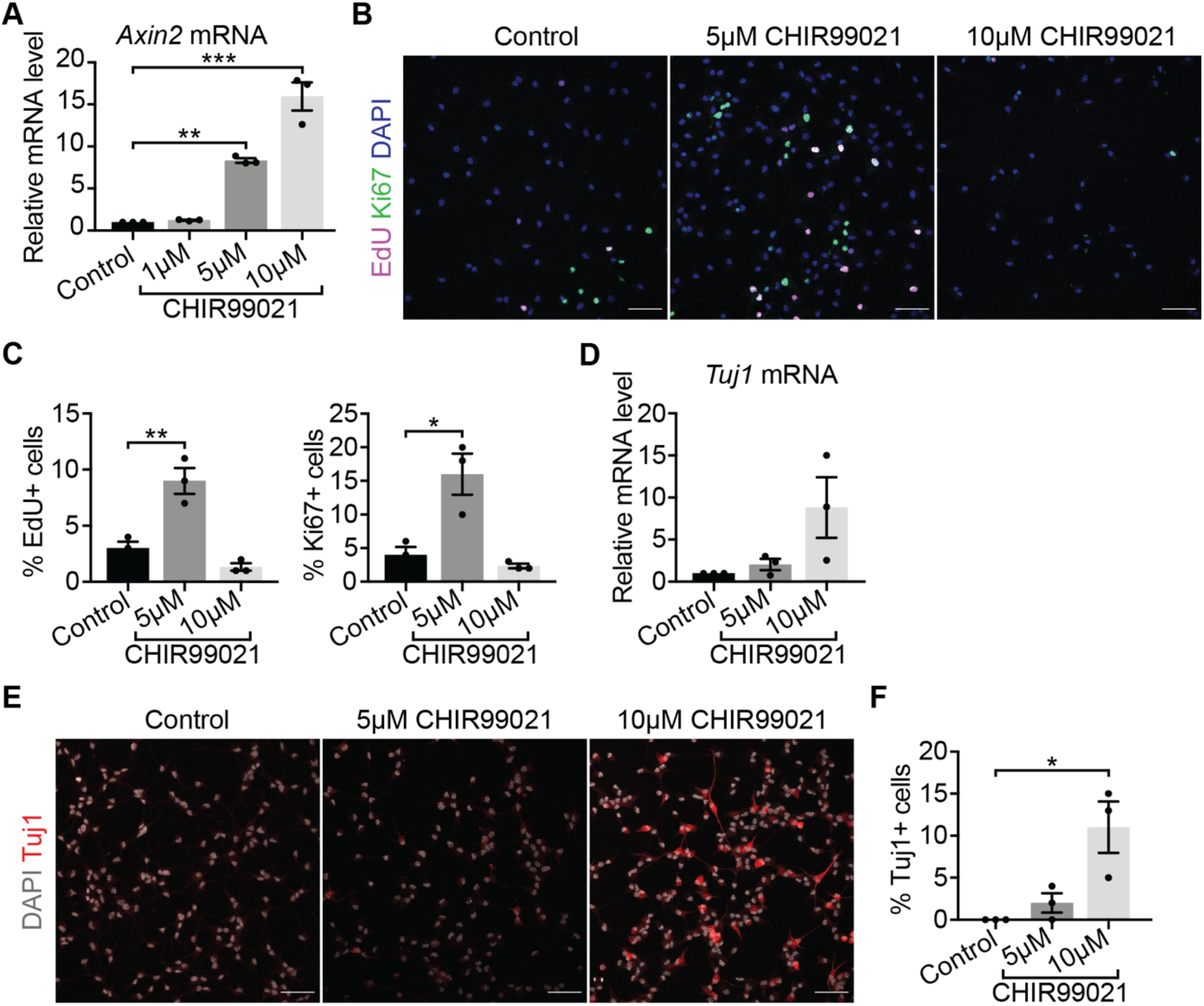
Stimulating Wnt/β-catenin signalling has dose-dependent effects in quiescent NSCs. (**A**) Upregulation of *Axin2* expression in quiescent NSCs treated with 5μM and 10μM CHIR99021 for 48hrs confirms stimulation of Wnt/β-catenin signalling. n=3. (**B**) Immunolabelling of the proliferation markers EdU and Ki67 and DAPI in quiescent NSCs treated with CHIR99021 compared with Control. Scale bars, 50μm. (**C**) Quantifications of the data shown in (**B**) of the proportion of EdU+ cells (Control, 3 ± 0.58%. 5μM CHIR99021, 9 ± 1.16%. 10μM CHIR99021, 1.33 ± 0.33%) and proportion of Ki67+ cells (Control, 4 ± 1.16%. 5μM CHIR99021, 16 ± 3.06%. 10μM CHIR99021, 2.33 ± 0.33%). Proliferation is increased by 5μM CHIR99021 treatment of quiescent NSCs. n=3. (**D**) *β-III-Tubulin (Tuj1)* expression is upregulated in quiescent NSCs treated with 10μM CHIR99021 for 48hrs compared with control (10μM CHIR99021, p=0.07). n=3. (**E**) Immunolabelling of Tuj1 and DAPI in quiescent NSCs treated with CHIR99021 compared with Control. Scale bars, 50μm. (**F**) Quantifications of the data shown in (**E**) of the proportion of Tuj1+ cells (Control, 0 ± 0%. 5μM CHIR99021, 2 ± 1.16%. 10μM CHIR99021, 11 ± 3.06%). 10μM CHIR99021 treatment of quiescent NSCs increases the proportion of Tuj1+ neurons. n=3. Statistics: Repeated measures one-way ANOVA with Dunnett’s multiple comparison test (**A, C, D** and **F**). (ns, p>0.05. *, p<0.05. **, p<0.01. ***, p<0.001). Error bars represent mean with SEM.

We then examined the effects of stimulating Wnt/β-catenin signalling on the proliferation and differentiation of quiescent NSCs. Treating quiescent NSCs with the highest concentration of CHIR99021 (10μM) did not significantly reduce the already very low levels of proliferation of quiescent NSCs (Fig. 6B, C). However, 10μM CHIR99021 treatment induced their neuronal differentiation, as seen by the increase in *Tuj1* expression and percentage of Tuj1+ cells (Fig. 6D-F), which resembles the effects we observed upon adding CHIR99021 to active NSCs (Fig. 5D-F). Remarkably, treating quiescent NSCs with 5μM CHIR99021 resulted in increases in the proportions of EdU-positive and Ki67-positive cells compared with vehicle-treated controls (Fig. 6B, C). This contrasts with the marked decrease in proliferation observed with 5μM CHIR99021 treatment of active NSCs (Fig. 5B-C). We also investigated neuronal differentiation and found that neither the expression levels of *Tuj1* nor the percentage of Tuj1+ cells were significantly changed in 5μM CHIR99021-treated quiescent NSCs relative to vehicle-treated control cells (Fig. 6D-F). Overall, these data show that moderate stimulation of Wnt/β-catenin signalling promotes the activation of quiescent NSCs while a higher level of stimulation promotes neuronal differentiation. This suggests that quiescent NSCs respond to Wnt/β-catenin signalling in a dose-dependent manner.

We next investigated whether the duration of CHIR99021 treatment also influences the response of quiescent NSCs. For this we performed a time course, where quiescent NSCs were treated with 5μM CHIR99021 for 24hrs, 48hrs and 72hrs. 5μM CHIR99021 treatment induced a similar increase in *Axin2* expression levels across all time points (Fig. 7A), indicating a constant level of Wnt/β-catenin stimulation over time. The proportion of Ki67-positive cells increased with time in 5μM CHIR99021-treated quiescent NSCs (Fig. 7B, C), while the expression of *Tuj1* was not significantly upregulated by sustained 5μM CHIR99021 treatment at any time point (Fig. 7D). Overall, this suggests that sustained 5μM CHIR99021 treatment promotes the activation of an increasing proportion of quiescent NSCs but does not initiate their neuronal differentiation.

**Figure 7:**
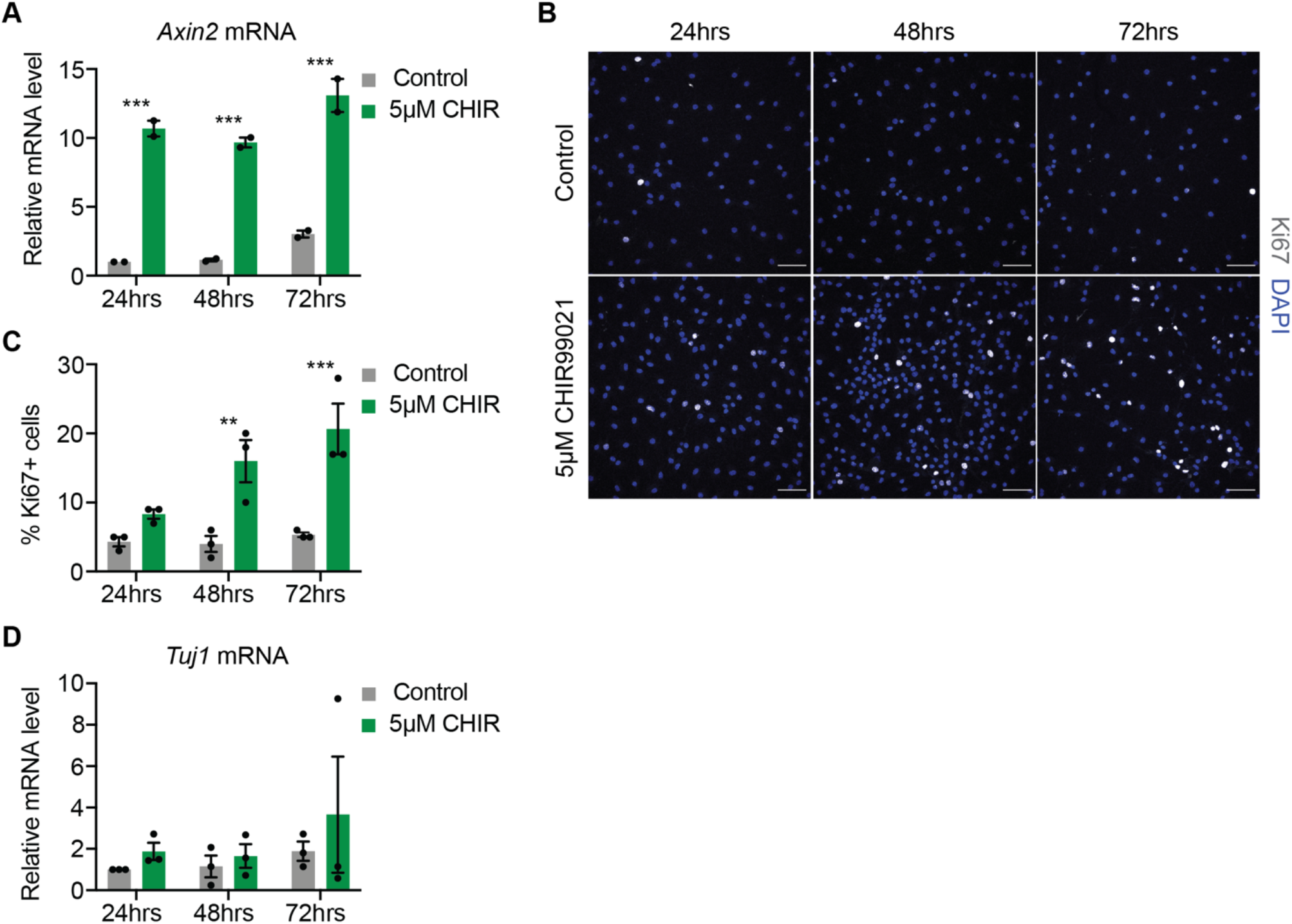
Quiescent NSC activation is sustained by prolonged Wnt/β-catenin stimulation. (**A**) Upregulation of *Axin2* is sustained by 5μM CHIR99021 treatment of quiescent NSCs for 24hrs, 48hrs and 72hrs. n=2. (**B**) Immunolabelling of the proliferation marker Ki67 and DAPI in quiescent NSCs treated with 5μM CHIR99021 for 24hrs, 48hrs and 72hrs. Scale bars, 50μm. (**C**) Quantification of the proportion of Ki67+ cells in (**B**). Proliferation increases with longer 5μM CHIR99021 treatments of quiescent NSCs (Control vs 5μM CHIR99021: 24hrs, 4.33 ± 0.67 vs 8.33 ± 0.67. 48hrs, 4 ± 1.16 vs 16 ± 3.06. 72hrs, 5.33 ± 0.33 vs 20.67 ± 3.67). n=3. (**D**) *Tuj1* expression is not significantly upregulated by 5μM CHIR99021 treatments of quiescent NSCs over time. n=3. Statistics: Two-way ANOVA with Sidak’s multiple comparisons test (**A, C** and **D**). (ns, p<0.05. **, p<0.01. ***, p<0.001). Error bars represent mean with SEM.

We confirmed the results obtained with the Wnt/β-catenin agonist CHIR99021 by treating quiescent NSCs with recombinant Wnt3a (rWnt3a). *Axin2* was upregulated in a dose-dependent manner by different concentrations of rWnt3a, with 500ng/ml rWnt3a inducing *Axin2* expression to a similar level as 5μM CHIR99021 (Fig. S5A). rWnt3a also increased Ki67 expression and EdU incorporation in a dose-dependent manner (Fig. S5B-D). These results confirm that directly stimulating Wnt/β-catenin signalling in quiescent NSCs promotes their activation.

## Discussion

In this study, we used genetic and pharmacological approaches to directly stimulate and inhibit Wnt/β-catenin signalling in quiescent and active NSCs, both *in vivo* and *in vitro*. We found that, whereas Wnt/β-catenin stimulation exerts dose-dependent and state-specific effects on NSCs, Wnt/β-catenin inhibition does not affect NSC homeostasis.

Our finding that baseline Wnt/β-catenin signalling is not essential for the maintenance of NSCs’ activation states or stem cell characteristics, differs from published reports showing that ablation of endogenous Wnt/β-catenin signalling affects NSC proliferation and multipotency (Wexler et al., 2009; Qu et al., 2013; Qu et al., 2010). Whilst we specifically targeted adult NSCs in our *in vivo* experiments, previous reports used knock-out mouse models, making it difficult to distinguish between developmental and adult phenotypes and between cell-autonomous and non-autonomous effects caused by the loss of Wnt/β-catenin signalling (Qu et al., 2010; Qu et al., 2013). Also, while we used genetic ablation of β-catenin in NSC cultures, other groups used approaches based on the extracellular sequestration of Wnt ligands (such as Fzd-CRD), which could affect both Wnt/β-catenin and non-canonical Wnt signalling (Wexler et al 2009 and Qu et al 2010). Non-canonical Wnt/PCP signalling maintains NSC quiescence in the V-SVZ but the role of Wnt/PCP in the SGZ has not yet been explored (Chavali et al., 2018). Our results corroborate work from Kuwabara et al., 2009 showing that loss of β-catenin in Sox2-positive proliferative progenitors *in vivo* does not affect the maintenance of the Sox2-positive progenitor pool. However, the authors used a retroviral approach to uniquely target proliferating cells and did not investigate the effects in NSCs, defined by their radial glial-like morphology to distinguish them from intermediate progenitors (Kuwabara et al., 2009). By conditionally targeting the deletion of β-catenin to Glast-positive NSCs, we directly assessed the role of Wnt/β-catenin signalling specifically in NSCs, including quiescent NSCs, and found that they are unaffected by the loss of β-catenin. Our results therefore showed that Wnt/β-catenin signalling is not required for NSC maintenance and function. We also found that loss of β-catenin in adult NSCs has no effect on the subsequent generation and proliferation of intermediate precursor cells or the generation of neuronally-committed precursors. This contradicts work from Kuwabara et al., 2009 who show that neuronal differentiation is impaired following the loss of β-catenin in Sox2-positive proliferative progenitors (Kuwabara et al., 2009). A possible explanation could be that Glast-dependent deletion occurs at a much earlier time point (when NSCs are still quiescent) compared with the acute β-catenin deletion in Sox2+ proliferative progenitors performed by Kuwabara and colleagues, therefore allowing more time for compensatory mechanisms. Of note, while we did not observe a difference in neurogenesis, we cannot exclude that β-catenin is needed for the maturation or function of newly generated neurons.

Numerous reports show that stimulating Wnt/β-catenin signalling increases adult neurogenesis by promoting progenitor proliferation and the maturation of newborn neurons (Lie et al., 2005; Jang et al., 2013; Seib et al., 2013; Qu et al., 2010). Our finding that Wnt/β-catenin signalling is dispensable for adult neurogenesis does not exclude a regulatory role of Wnt/β-catenin stimulation on NSCs and their progeny. To directly stimulate Wnt/β-catenin signalling in NSCs *in vivo* we first tried a genetic approach to stabilise β-catenin, which involved the deletion of exon 3, harbouring the GSK3β phosphorylation and a-catenin binding sites (Harada et al., 1999). Upon recombination, we observed increased Wnt/β-catenin activity in NSCs but also their displacement from the SGZ. Disruption of β-catenin binding to actin via a-catenin could affect the adhesion properties of recombined NSCs, contributing to the displacement phenotype we observed and to their subsequent loss. Indeed, disruption of β-catenin/cadherin signalling and loss of cell-cell contacts can disrupt tissue organisation and promote cell migration (Berx et al., 2001; Conacci-Sorrell et al., 2002; Taniguchi et al., 2006; Machon et al., 2007; Kadowaki et al., 2007). Interestingly, deletion of β-catenin did not appear to affect cell-cell contacts and NSC positioning. This could be due to the differences between inhibition of baseline Wnt/β-catenin signalling *versus* gain-of-function effects or by compensation from g-catenin, which has been shown to be able to replace β-catenin’s function at adherens junctions in hepatocytes (Wickline et al., 2013). The early appearance of the displacement phenotype prevented us from using this mouse model to investigate the effects of Wnt/β-catenin stimulation on NSC behaviour. To overcome this problem, we used an *in vitro* model of hippocampal NSCs, which allows the culture of pure and homogeneous populations of NSCs in a controlled environment. We could show that stimulating Wnt/β-catenin signalling promotes neuronal differentiation of active NSCs, whilst promoting the proliferation or differentiation of quiescent NSCs in a dose-dependent manner. The dose-dependent effects of stimulating Wnt/β-catenin signalling in quiescent NSCs shown here, could provide a possible explanation for some of the contradictions reported in the literature, as different techniques used to modulate Wnt could stimulate Wnt/β-catenin signalling levels to varying degrees (Lie et al., 2005; Jang et al., 2013; Seib et al., 2013; Qu et al., 2010). For example, hippocampal specific overexpression of Wnt ligands, which was found to promote neuronal differentiation (Lie et al., 2005), could induce a higher level of Wnt/β-catenin stimulation compared with knocking-out a Wnt inhibitor, which was shown to enhance progenitor proliferation *in vivo* (Seib et al., 2013; Jang et al., 2013). These non-targeted approaches could induce secondary or non-cell autonomous effects, therefore, novel *in vivo* tools will be needed to specifically modulate and compare stimulated Wnt/β-catenin levels in active and quiescent adult NSCs to confirm this hypothesis.

Dose-dependent effects of Wnt/β-catenin signalling have been reported in other systems, such as hematopoietic stem cells, intestinal stem cells, during cortical development and hippocampal cellular specification (Luis et al., 2011; Hirata et al., 2013; Machon et al., 2007). Specifically, low levels of Wnt/β-catenin stimulation preserve the self-renewal capacity of hematopoietic stem cells whereas higher Wnt/β-catenin dosages enhance myeloid lineage differentiation (Luis et al., 2011). In NSCs, one potential mechanism mediating this dose-dependent response could be the dual regulation of *NeuroD1* by Sox2 and the Wnt/β-catenin downstream effector TCF/LEF (Kuwabara et al., 2009). *NeuroD1* harbours overlapping binding sites for both Sox2 and TCF/LEF, where Sox2 binding represses *NeuroD1* expression in NSCs and Wnt/β-catenin signalling induces *NeuroD1* expression in a dose-dependent manner (Kuwabara et al., 2009). The higher level of Wnt/β-catenin stimulation induced by 10μM CHIR99021 could override the Sox2 repression of *NeuroD1* in quiescent NSCs to induce neuronal differentiation, whereas the lower 5μM CHIR99021 Wnt/β-catenin stimulation may not be sufficient. Whilst lower Wnt/β-catenin stimulation in quiescent NSCs promotes their activation, we also observed a mild upregulation of *Tuj1* (not significant). Further investigation is needed to determine if this activation is linked to a loss of stemness and increased neuronal commitment or if Wnt/β-catenin activated NSCs are able to self-renew and return to a resting state of quiescence.

Whilst our results showed that Wnt/β-catenin signalling is dispensable for NSC homeostasis in young adult mice, NSCs still respond to Wnt/β-catenin stimulation, which could be important to mediate their response to external stimuli. For example, Wnt ligand levels increase following exercise in the DG of aged mice (Okamoto et al., 2011) and expression of the granule neuron-derived Wnt inhibitor sFRP3, is decreased by granule neuron activity (Jang et al., 2013) suggesting that the level of Wnt/β-catenin signalling in the DG is regulated by extrinsic factors. Decreased Wnt/β-catenin signalling levels with age contribute to age-related decline in adult hippocampal neurogenesis (Miranda et al., 2012; Seib et al., 2013; Okamoto et al., 2011) and Wnt-mediated adult hippocampal neurogenesis has been shown to be important for learning and memory (Jessberger et al., 2009). This suggests that Wnt/β-catenin signalling integration in NSCs could be an important regulator of external stimuli on adult hippocampal neurogenesis and hippocampal functions.

Our finding that quiescent and active NSCs are differentially responsive to Wnt/β-catenin signalling means that quiescent and active NSCs could coordinate their response to stimulated Wnt/β-catenin signalling *in vivo* to simultaneously promote neurogenesis and NSC activation. These state-specific responses to Wnt/β-catenin signalling could be driven by quiescent and active NSCs integrating different niche-derived signalling pathways. For example, Notch and BMP signalling are higher in quiescent NSCs compared with active NSCs, where they maintain and promote quiescence (Lavado et al., 2014; Ables et al., 2010; Imayoshi et al., 2010; Mira et al., 2010; Bonaguidi et al., 2008). Wnt/β-catenin signalling cooperatively regulates many developmental processes together with Notch and BMP signalling (reviewed by (Muñoz Descalzo et al., 2012; Itasaki et al., 2010)). Therefore, Wnt/β-catenin signalling levels modulated by external stimuli could play a key role in determining the response of quiescent NSCs to other niche cues.

In conclusion, our results show that Wnt/β-catenin signalling is dispensable for NSC homeostasis but also that Wnt/β-catenin stimulation exerts dose-dependent and state-specific effects on NSCs that could contribute to the regulaton of adult hippocampal neurogenesis in response to external stimuli. Further work is needed to understand the downstream molecular mechansisms by which levels of Wnt/β-catenin signalling induce differential effects across various cell types.

## Acknowledgments

We gratefully acknowledge Maria del Mar Masdeu for technical advice and training; Lan Chen for technical support; Rekha Subramaniam, Nicholas Chisholm and the Crick BRF staff for managing the mouse colonies; Donald Bell from the Crick Advanced Light Miscroscopy science technology platform for training and technical advice; Lotte Carr from the Crick Flow Cytometry science technology platform for technical support and assistance and members of the Guillemot lab for their discussions and support.

## Additional Information

### Author Contributions

Conceptualisation [SA, FG, NU]; Data curation [SA]; Formal analysis [SA]; Validation [SA, LH]; Resources [LH]; Investigation [SA, LH, OP]; Visualisation [SA, LH, OP, PR]; Methodology [SA, LH]; Writing-original draft [SA, FG, NU], Project administration [FG]; Supervision [FG, NU], Funding acquisition [FG, NU].

## Materials and Methods

Contact for reagent and resource sharing: François Guillemot

## Experimental model and subject details

### Mouse models

All procedures involving animals and their husbandry were performed according to the guidelines of the Francis Crick Institute, national guidelines and laws. This study was approved by the Animal Ethics Committee and by the UK Home Office (PPL PB04755CC). Mice were housed in standard cages under a 12hr light/dark cycle with ad libitum access to food and water. All experimental mice were of mixed background. Founder mice were crossed with MF1 mice, and then backcrossed to littermates of the F1 generation. *Glast-CreERT2 (Slc1a3^tm(cre/ERT2)Mgoe^)* (Mori et al., 2006) mice were crossed with *Rosa26-floxed-stop-YFP (RYFP; Gt(ROSA)26Sor^tm1(EYFP)Cos^)* (Srinivas et al., 2001) mice to generate NSC specific tamoxifen inducible mice with a YFP recombination reporter. These mice were then crossed with the experimental strains: β-cat^fl/fl ex3-6^ *(Ctnnb1^tm2Bir^)* (Huelsken et al., 2001), β-cat^fl/fl ex2-6^ *(Ctnnb1^tm2Kem^)* (Brault et al., 2001) and Catnb^lox(ex3)^ *(Ctnnb1^tm1Mmt^)* (Harada et al., 1999). *Glast-CreERT2; β-cat^fl/fl ex3-6^; RYFP mice, Glast-CreERT2; β-cal^fl/fl ex2-6^; RYFP* mice and *Glast-CreERT2; Catnb^lox(ex3)^; RYFP* mice were crossed with BATGAL Wnt/β-catenin reporter mice (Maretto et al., 2003) to introduce the *BATGAL* Wnt/β-catenin reporter allele.

Experimental groups included a combination of mice from different litters within each strain and both males and females were used for all *in vivo* studies.

### Primary cell cultures

Adult hippocampal NSC cultures were derived as previously described (Blomfield et al., 2019). Briefly, 7-8 week old mice were sacrificed and the DG dissected (previously described by (Walker et al., 2014). For each new line, the two DGs of a single mouse of the desired genotype were dissociated using the Neural Tissue dissociation kit (P) (Milteny Biotec, 130-092-628). NSCs were amplified in non-adherent conditions as neurospheres in basal media (DMEM:F12 + L-Glutamine + sodium bicarbonate (Gibco, 11320)) + 1mg/ml KCl (Sigma, P5405) + 2mg/ml BSA (Sigma, A9056)) + 1x Neurocult Supplement (Stem Cell Technologies, 05701) + 1x Penicillin/Streptomycin (ThermoFischer Scientific, 15140) + 10ng/ml FGF2 (Protech, 450-33) + 20ng/ml EGF (Protech, 315-09) + 2μg/ml Heparin (Sigma, H3393) for at least two passages before dissociation to adherent cultures. NSCs were propagated in basal media (DMEM/F-12 + Glutamax (Invitrogen 31331-093)) + 1x N2 Supplement (R&D Systems, AR009) + 1x Penicillin/Streptomycin (ThermoFischer Scientific, 15140) + 2μg/ml Laminin (Sigma, L2020) + 5μg/ml Heparin (Sigma, H3393) + 20ng/ml FGF2 (Protech, 450-33). Neurospheres and adherent NSCs were incubated 37°C, 5% CO_2_.

## Method details

### Tamoxifen administration

To induce activation of CreERT2 recombinase, two-month old *GlastCreERT2; β-cat^fl/fl ex3-6^; RYFP* mice were administered 2mg (57-67mg/kg) 4-hydroxy-tamoxifen (Sigma, H6278) by ip injection at the same time each day for five consecutive days. Due to high numbers of deaths, and following veterinary advice, we changed to use tamoxifen (Sigma, T5648) for all other experiments. Two-month-old mice were administered 2mg (57-67mg/kg) tamoxifen (Sigma, T5648) by intraperitoneal (ip) injection at the same time each day for five consecutive days.

Approximately 2 weeks after tamoxifen administration, *Glast-CreERT2; Catnb^lox(ex3)^; RYFP* mice developed a skin phenotype due to the stabilisation of β-catenin in Glast-expressing activated hair follicle stem and progenitor cells (Reichenbach et al., 2018). We were able to collect a sufficient number of samples from this first cohort of mice to perform meaningful analysis 30 days after tamoxifen injections. However, all subsequent analyses were performed a maximum of 10 days after tamoxifen injections, following veterinary advice, before the skin phenotype developed.

### Tissue preparation and immunofluorescence

Mice were transcardially perfused, under terminal anaesthesia, with phosphate-buffered saline (PBS) for 2mins, followed by 4% paraformaldehyde (PFA) in PBS for 12mins. Brains were removed and post-fixed overnight in 4% PFA at 4°C with rocking and then transferred to PBS containing 0.02% Sodium Azide. Brains were coronally sectioned at a thickness of 40μm using a vibratome (Leica).

Cultured NSCs were fixed with 4%PFA in PBS for 10mins at room temperature and then washed with PBS.

Antigen retrieval was performed in brain sections prior to immunofluorescence using the following antibodies: Ki67 (BD Biosciences, 550609), Sox2 (EBioscience, 14-9811-82) and Tbr2 (Abcam, ab183991). For antigen retrieval, samples were incubated at 95°C for 2min in sodium citrate buffer (10mM, pH6.0). Following antigen retrieval, samples were processed for immunofluorescence as previously described (Blomfield et al., 2019). Prior to mounting brain sections were incubated with 2μg/ml DAPI (Sigma, D9542) in 1:1 PBS:H_2_O for 30min and fixed NSCs were incubated with 2μg/ml DAPI (Sigma, D9542) in 1:1 PBS:H_2_O for 10min at room temperature. Primary and secondary antibodies and dilutions are listed in Supplementary Table 1.

EdU was detected prior to DAPI incubation using the Click-iT™ EdU Alexa Fluor^®^ 647 detection kit (Invitrogen, C10340) following manufacturer’s instructions.

### Microscopic analysis

Immunofluorescence samples were imaged using a SP5 confocal microscope (Leica) with a 40X oil objective lens (Leica). For fixed NSCs, three random regions of each coverslip were imaged with a 1μm z-step. For brain sections both left and right DG of every twelfth 40μm section along the rostro-caudal axis were imaged, with a 1μm z-step through the whole 40μm section.

### Cell treatments, constructs, transfection and transduction

To induce quiescence, cells were plated in flasks or on coverslips in active culture conditions and allowed to adhere overnight. The next day, media was replaced with basal media plus 20ng/ml BMP4 (R&D Systems, 5020-BP) and incubated for 72hrs at 37°C, 5% CO_2_ to establish quiescence as previously described (Blomfield et al., 2019). All experiments using quiescent NSCs were started following this 72hrs incubation with BMP4 to ensure robust quiescence induction.

To label cells in S-phase of the cell cycle, EdU (Invitrogen, C10340) was added to the media of cultured cells at a final concentration of 10μM 1hr before fixation.

To delete β-catenin in NSCs derived from β-cat^fl/fl ex3-6^; RYFP mice, active and quiescent β-cat^fl/fl ex3-6^ NSCs were transduced with either adenovirus control (Ad-CMV-Null, Vector Biolabs, 1300) or adenovirus expressing Cre (Ad-CMV-iCre, Vector Biolabs, 1045) at a MOI=100. Samples of active β-cat^fl/fl ex3-6^ NSCs were collected 1-, 2- and 3-days after transduction for qPCR, immunofluorescence and Western blot analyses. Samples of quiescent β-cat^fl/fl ex3-6^ NSCs were collected 2-, 4- and 6-days after transduction for qPCR and immunofluorescence analyses. To stimulate Wnt/β-catenin signalling in active β-cat^fl/fl ex3-6^ NSCs, 48hrs after transduction, media was replaced with basal media plus 5μM CHIR99021 (BioVision, 1677-5) or DMSO vehicle control and incubated for 24hrs at 37°C, 5% CO_2_ before collection in TRIzol for RNA extraction, cDNA synthesis and qPCR analysis. To determine the reactivation rate of quiescent β-cat^fl/fl ex3-6^ NSCs, 48hrs after transduction, media was replaced with basal media and samples were collected 3- and 6-days later for immunofluorescence analysis. Samples were also collected in basal media plus 20ng/ml BMP4, 48hrs after transduction for the 0hrs time point. Media was refreshed every 72hrs. To determine how chronic loss of β-catenin affects neurogenesis and gliogenesis, at 6-, 12- and 18-days after transduction, the media on transduced active β-cat^fl/fl ex3-6^ NSCs was replaced with basal media minus 20ng/ml FGF2 plus 2% B27 to promote neuronal differentiation and basal media minus 20ng/ml FGF2 plus 2% FBS to promote astrocytic differentiation. Cells were incubated in these differentiation media condition for 5 days at 37°C 5% CO_2_ before fixation.

To stimulate Wnt/β-catenin signalling in active NSCs, media on cells was replaced with basal media supplemented with either DMSO vehicle control, 1μM CHIR99021, 5μM CHIR99021 or 10μM CHIR99021 and incubated for 48hrs at 37°C 5% CO_2_ before sample collection. To stimulate Wnt/β-catenin signalling in quiescent NSCs, media on cells was replaced with basal media plus 20ng/ml BMP4 supplemented with either DMSO vehicle control, 1μM CHIR99021, 5μM CHIR99021 or 10μM CHIR99021 and incubated for 24hrs, 48hrs and 72hrs at 37°C 5% CO_2_ before sample collection. To compare the effects of CHIR99021 treatment vs rWnt3a (R&D Systems, 1324-WN) in quiescent NSCs, media on cells was replaced with basal media plus 20ng/ml BMP4 supplemented with either DMSO (vehicle control for CHIR99021), 5μM CHIR99021, 0.1%BSA/PBS (vehicle control for rWnt3a), 100ng/ml rWnt3a, 250ng/ml rWnt3a or 500ng/ml rWnt3a and incubated 48hrs at 37°C 5% CO_2_ before sample collection. Media was refreshed every 24hrs.

### Protein purification and Western blots

For Western blot analysis, samples were washed with ice-cold PBS and scraped in Lysis buffer (ThermoFischer Scientific, 87788) supplemented with phosphatase inhibitor (ThermoFischer Scientific, 78420), EDTA (ThermoFischer Scientific, 87788) and protease inhibitor (ThermoFischer Scientific, 87786). Cells were lysed at 4°C for 20mins under rotation and then centrifuged at 13,000 rpm at 4°C for 20mins. The supernatant was transferred to a clean Eppendorf and an equal volume of Laemmli sample buffer (Sigma, S3401-10VL) was added before boiling at 95°C for 5mins.

Samples were run on a polyacrylamide gel at 120V and then transferred onto a nitrocellulose membrane. Membranes were blocked with 5% milk in TBS-Tween or 5% BSA in TBS-Tween before incubation with primary antibody diluted in 5% milk in TBS-Tween (Actin, β-catenin and Vinculin) or 5% BSA in TBS-Tween + 0.02% sodium azide (Wnt7a) overnight at 4°C under rotation. The following day, membranes were washed with TBS-Tween and incubated with secondary antibody diluted in 5% milk in TBS-Tween, for 2hrs at room temperature with rocking. Detection was performed using ECL Western Blotting reagents (Sigma, GERPN2106) according to manufacturer’s instructions and blots were developed using Kodak films. Primary and secondary antibodes and dilutions are listed in Supplementary Table 1.

### Dentate gyrus dissections and fluorescence activated cell sorting (FACS)

Two-month-old Control, β-cat^del ex3-6^ and β-cat^del ex2-6^ mice were injected with tamoxifen for 5 consecutive days and culled ten days later. The dentate gyrus was dissected as previously described (Walker et al., 2014). Dissected dentate gyri were dissociated using the Neural Tissue dissociation kit (P) (Milteny Biotec, 130-092-628) as described (Harris et al., 2020). Dissociated cells were centrifuged and resuspended in 750μl recovery media (0.5% PBS/BSA in DMEM/F12 without phenol red (Sigma, DG434) and 1μg/ml DAPI). Cells were sorted on a MoFLo XDP cell sorter machine (Beckman Coulter) using a 100μm nozzle at a maximum sort speed of 5000 events per second and an efficiency of more than 80%. Events were first gated to remove debris (fsc-h vs ssc-h), aggregates (pulse width vs area then pulse-height vs area/width) and dead cells (fsc-h vs DAPI fluorescence). Cells were then gated for YFP expression using a YFP-negative control mouse. Mice were processed for FACS separately and YFP-positive cells from individual samples were collected directly in 700μl RL lysis buffer from the RNeasy^®^ Mini Kit (Qiagen, 74104) for RNA extraction.

### RNA extraction, cDNA synthesis and qPCR

For FACS experiments, cells were lysed using the RL lysis buffer from the RNeasy^®^ Mini Kit (Qiagen, 74104). For all other experiments, cells were lysed using TRIzol reagent (Ambion, 15596018). RNA was extracted using the RNeasy® Mini Kit (Qiagen, 74104) or the Direct-zol™ RNA MiniPrep Kit (Zymo Research, R2052) according to manufacturer’s instructions. cDNA was synthesised using the Maxima First Strand cDNA Synthesis Kit (ThermoScientific, K1641) or the High Capacity cDNA Reverse Transcription Kit (Applied Biosystems, 4387406) according to manufacturer’s instructions. Gene expression levels were measured using TaqMan Gene expression assays (Applied Biosystems, Supplementary Table 2) and qPCR was performed on a QuantStudio Real-Time PCR machine (ThermoFisher). Gene expression was calculated relative to the endogenous controls *GAPDH* and *ActinB* (dCt), and to the control samples to provide a delta delta cycle threshold (ddCt) value. ddCT values were then coverted to fold change of gene expression relative to control. All samples were measured in technical duplicates for each qPCR run and averaged.

### RNA Sequencing and Analysis

The publicly available single cell RNA sequencing dataset of *in vivo* NSCs (Harris et al., 2020) was accessed from the Gene Expresssion Omnibus (GSE159768) with associated code (https://github.com/harrislachlan/lifelong_stemcells). The final dataset contained 2,947 which were classified as dormant, resting and active NSCs as described by (Harris et al., 2020). For this study, the metadata was appended to classify dormant and resting NSCs as quiescent NSCs. Actie NSC classification remained the same. Differential gene expression analysis was performed using the FindMarkers function in Seurat (Version 3.1.1) (Stuart et al., 2019) using the Pearson residuals located in the “scale.data” slot of the SCT assay using Student’t t-test (Hafemeister et al., 2019). All genes that were expressed by a minium of 10% of cells were tested for differential expression. No minimum log fold change threshold was enforced. The heatmap in Fig. 1A was generated using the DoHeatmap function.

The publicly available bulk RNA sequencing of active and quiescent NSCs cultures ((Blomfield et al., 2019) was accessed from the Gene Expresssion Omnibus (GSE116997).

## Quantifications and statistical analysis

### Immunofluorescence quantifications

NSCs in DG images from BATGAL mice were identified as having a DAPI-positive cell body located within the subgranular zone (SGZ) co-localised with a GFAP-positive radial projection through the granule cell layer (GCL) spanning a minimum of 2 cell nuclei away from the NSC nucleus. NSCs in all other DG images were identified as above but also as having a Sox2-positive cell body. Displaced NSCs were identified as above but with the DAPI-positive Sox2-positive cell body being located more than 2 nuclei’s distance away from the SGZ. To measure the intensity of BATGAL immunofluorescence staining, the nucleus of each BATGAL+ NSC was manually outlined according to DAPI, and the mean pixel value of the channel of interest was measured using FIJI software. Each value was normalised to the average BATGAL intensity level measured in DAPI-positive cells in the GCL from the same z-plane as each NSC. For quantifying the proportion of cells expressing specific markers in DG images, a minimum of 100 cells were counted per animal for a minimum of 3 animals per genotype. For quantifying the total number of cells/SGZ length mm, a minimum of 2 sections per series per mouse was quantified for a minimum of 3 mice per genotype. The freehand line tool in Fiji (Schindelin et al., 2012) was used to measure the length of the SGZ in μm.

For quantification of cultured NSCs immunolabelled samples, the DAPI channel was used to generate a mask of all nuclei using the Fiji software. This mask was used to measure the average pixel intensity for each channel within the area of each nuclei. The proportion of cultured NSCs expressing specific markers was quantified by setting a signal threshold and expressing the number of positive cells as a percentage of the total number of cells. For each experiment, at least 100 cells were quantified across 3 biological replicates unless stated otherwise.

### Statistical analysis

All statistical analyses were performed using the GraphPad Prism seven software (RRID:SCR_002798). Statistical analyses were conducted using a two-tailed unpaired Student’s t-test; two-tailed paired Student’s t-test; Ordinary one-way ANOVA with Tukey’s multiple comparison test; Repeated measures one-way ANOVA with Dunnett’s multiple comparison test and two-way ANOVA with Sidak’s multiple comparisons tests as appropriate. All error bars represent the mean ± SEM. Significance is stated as follows: ns, p>0.05. *, p<0.05. **, p<0.01. ***, p<0.001. The statistical details and sample sizes for each experiment are recorded in the Figure legends.

---

**Supplementary Table 1:** Primary and secondary antibodies.

| Target Molecule | Species | Procedure | Dilution | Supplier | Catalogue # |
| --- | --- | --- | --- | --- | --- |
| Actin | Rabbit | WB | 1:1000 | Sigma-Aldrich | A2066 |
| β-catenin | Mouse | IF-*in vivo* | 1:100 | BD Biosciences | 610154 |
|  |  | IF-*in vitro* | 1:250 |  |  |
|  |  | WB | 1:2000 |  |  |
| β-galactosidase | Chicken | IF | 1:1000 | Aves Lab Inc. | BGL-1010 |
| CyclinD1 | Rabbit | IF | 1:25 | ThermoScientific | RM-9104 |
| DCX | Goat | IF | 1:50 | Santa-Cruz | Sc-8066<br>(Discontinued) |
| GFAP | Rat | IF | 1:800 | Invitrogen | 13-0300 |
| GFP | Chicken | IF | 1:2000 | Abcam | ab13970 |
| Ki67 | Mouse | IF | 1:100 | BD Biosciences | 550609 |
| Map2 | Mouse | IF | 1:200 | Sigma | M4403 |
| mCherry | Rabbit | IF | 1:500 | GeneTex | GTX128508 |
| Sox2 | Rat | IF | 1:400 | EBioscience | 14-9811-82 |
| Tbr2 | Rabbit | IF | 1:200 | Abcam | ab183991 |
| Tuj1 | Rabbit | IF | 1:400 | Covance | PRB-435P |
| Tuj1 | Mouse | IF | 1:400 | Covance | MMS-435P |
| Wnt7a | Rabbit | WB | 1:500 | Abcam | ab100792 |
| Vimentin | Mouse | WB | 1:2000 | Sigma | V9131 |
| Chicken IgG | Donkey | IF-488 | 1:500 | Jackson | 703-545-155 |
| Mouse IgG | Donkey | IF-488 | 1:500 | Jackson | 715-546-151 |
| Rat IgG | Donkey | IF-Cy3 | 1:500 | Jackson | 712-166-153 |
| Rabbit IgG | Donkey | IF-Cy3 | 1:500 | Jackson | 711-166-152 |
| Mouse IgG | Donkey | IF-Cy3 | 1:500 | Jackson | 715-166-151 |
| Goat IgG | Donkey | IF-647 | 1:500 | Jackson | 705-605-147 |
| Mouse IgG | Donkey | IF-647 | 1:500 | Jackson | 715-606-151 |
| Rabbit IgG | Donkey | IF-647 | 1:500 | Jackson | 711-606-152 |
| Rat IgG | Donkey | IF-647 | 1:500 | Jackson | 112-175-167 |
| Mouse IgG | Rabbit | WB-HRP | 1:1000 | Dako | P0161 |
| Rabbit IgG | Goat | WB-HRP | 1:1000 | Dako | P0448 |

**Supplementary Table 2:** List of TaqMan probes from Applied Biosystems used for qPCR gene expression assays.

| Gene | Assay ID | Catalogue Number |
| --- | --- | --- |
| ActinB | Mm00607939_s1 | 4352933E |
| Ascl1 | Mm03058063_m1 | 4331182 |
| Axin2 | Mm00443610_m1 | 4331182 |
| GAPDH | Mm99999915_g1 | 4352932E |
| HopX | Mm00558630_m1 | 4331182 |
| Id4 | Mm00499701_m1 | 4331182 |
| Nestin | Mm00450205_m1 | 4331182 |
| Ngn2 | Mm00437603_g1 | 4331182 |
| Sox2 | Mm00488369_s1 | 4331182 |
| Tubb3 (Tuj1) | Mm00727586_s1 | 4331182 |
| β-catenin exons 4-5 | Mm01350386_g1 | 4351372 |

## Data and Code Availability

No new data was generated for this study. Code will be deposited in Github upon publication.

## Supplementary Figures

**Figure S1:**
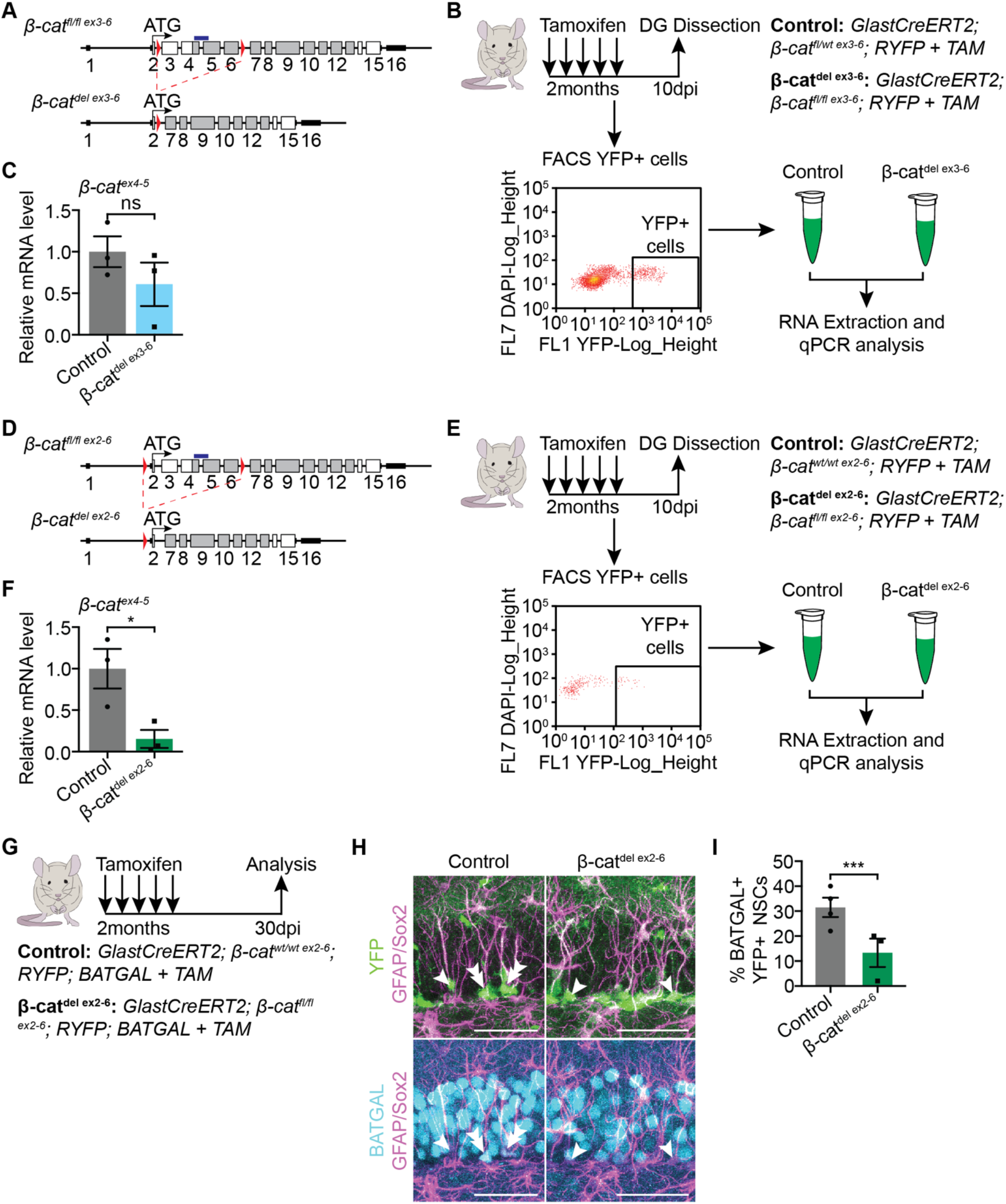
Analysis of *β-catenin* recombination in β-cat^del ex3-6^ and β-cat^del ex2-6^ mice and successful inhibition of Wnt/β-catenin signalling in β-cat^del ex2-6^ mice. (**A**) *β-cat^fl/fl ex3-6^* allele before and after recombination. LoxP sites (red arrows) flank exons 3-6, which encode the N-terminal domain and armadillo repeats 1-4. Grey boxes indicate armadillo repeats. Blue box indicates the sequence recognised by the TaqMan probe used for qPCR analysis in (**C**). (**B**) Two-month-old Control and β-cat^del ex3-6^ mice were injected with tamoxifen once daily for 5 consecutive days to induce recombination of the *β-cat^fl/fl ex3-6^* allele. Ten days after the first tamoxifen injection, the DG of individual mice were dissected and YFP+ cells were collected by flow cytometry for RNA extraction and qPCR analysis. (**C**) *β-cat^ex4-5^*(blue box, **A**) expression is not significantly different between YFP+ cells collected by flow cytometry from the DG of Control and β-cat^del ex3-6^ mice indicating failed recombination. n=3. (**D**) *β-cat^fl/fl ex2-6^* allele before and after recombination, as in (**A**). **(E)** Two-month-old Control and β-cat^del ex3-6^ mice were injected with tamoxifen once daily for 5 consecutive days. Ten days after the first tamoxifen injection, the DG of individual mice were dissected and YFP+ cells were collected by flow cytometry for RNA extraction and qPCR analysis. **(F)** *β-cat*(blue box, **D**) expression is significantly decreased in β-cat^del ex2-6^ YFP+ cells compared with Control YFP+ cells indicating successful *β-cat^fl/fl ex2-6^* recombination. n=3. (**G**) Two-month-old Control and β-cat^del ex2-6^ mice, crossed with BATGAL Wnt/β-catenin reporter mice, were administered tamoxifen for 5 consecutive days and sacrificed 30 days after the first tamoxifen injection. (**H**) BATGAL (β-galactosidase), GFAP and Sox2 immunolabelling in the DG of Control and β-cat^del ex2-6^ mice crossed with *BATGAL* mice. Arrowheads indicate BATGAL-NSCs and double arrowheads indicate BATGAL+ NSCs. Scale bar, 50μm. (**I**) Quantification of the proportion of BATGAL+ NSCs in **(H)**. The decreased proportion of BATGAL+ NSCs in β-cat^del ex2-6^ mice (13.33 ± 5.69%, n=3) compared with Control (31.5 ± 3.88%, n=4) indicates successful inhibition of Wnt/β-catenin signalling following recombination of the *β-cat^fl/fl ex2-6^* allele. Statistics: unpaired Student’s t-test **(C, F** and **I**). (ns, p>0.05. *, p<0.05. ***, p<0.001). Error bars represent mean with SEM.

**Figure S2:**
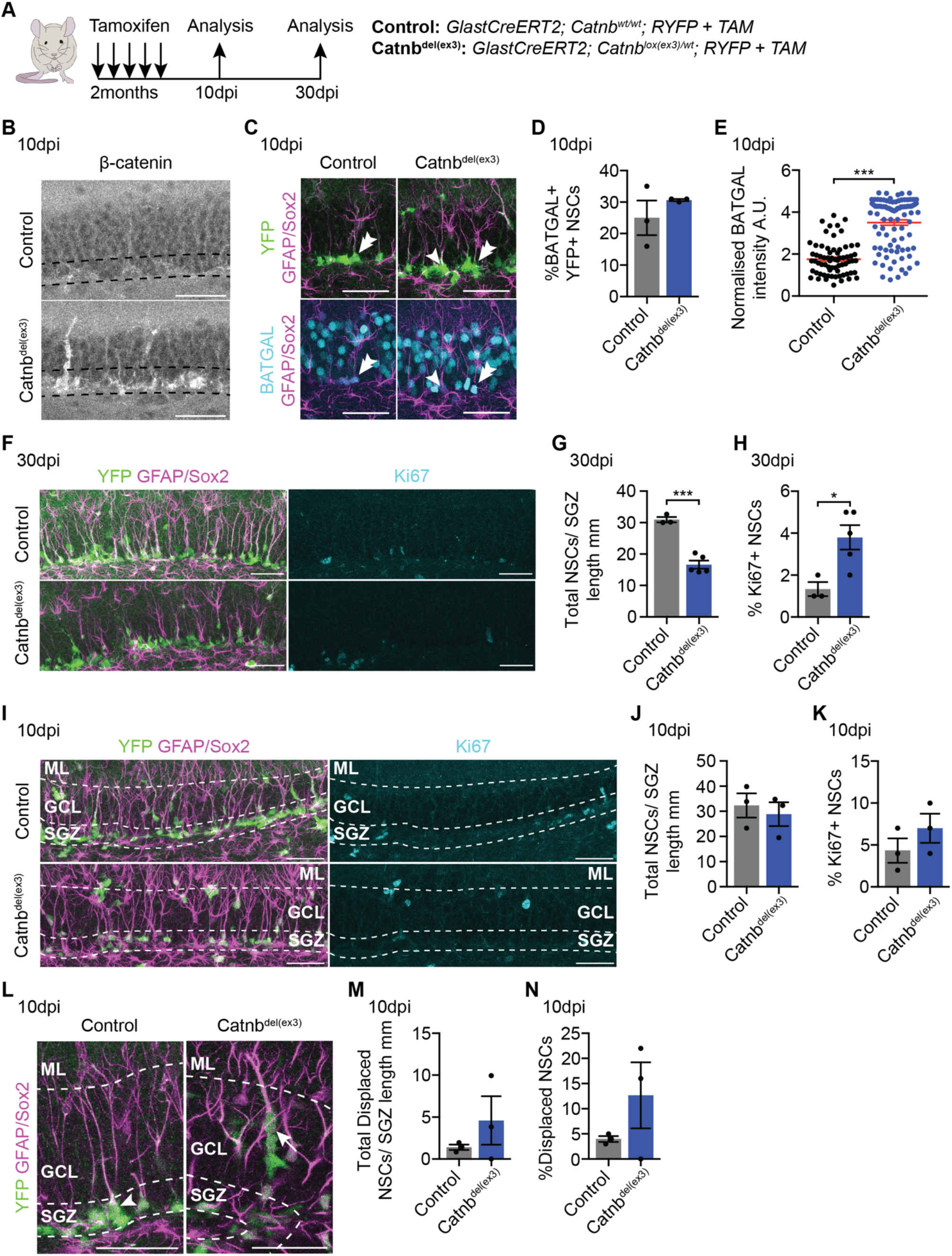
Stabilising β-catenin in NSCs *in vivo* displaces them from their correct niche location. (**A**) Two-month-old Control and Catnb^del(ex3)^ mice were administered tamoxifen for 5 consecutive days and sacrificed 10 and 30 days after the first tamoxifen injection. (**B**) β-catenin immunolabelling in the DG of Control and Catnb^del(ex3)^ mice 10 days after tamoxifen. Increased β-catenin staining in Catnb^del(ex3)^ mice indicates successful recombination. Dashed lines mark the SGZ. Scale bars, 50μm. (**C**) BATGAL (β-galactosidase), GFAP and Sox2 immunolabelling in the DG of Control and Catnb^del(ex3)^ mice crossed with BATGAL mice, 10 days after tamoxifen. Arrowheads indicate BATGAL-NSCs and double arrowheads indicate BATGAL+ NSCs. Scale bar, 50μm. (**D**) Quantification of the proportion of BATGAL+ NSCs in (**C**). The proportion of BATGAL+ NSCs is unchanged between Control (25 ± 5.51%) and Catnb^del(ex3)^ (30.67 ± 0.33%) 10 days after tamoxifen, which corresponds to the proportion of NSCs expressing *β-catenin* by RNA sequencing (32% quiescent NSCs. n=3. (**E**) Quantification of the data shown in (**C**) of the intensity of BATGAL staining in BATGAL+ NSCs normalised to the BATGAL intensity in neighbouring DAPI+ GCL cells. BATGAL reporter intensity is increased in BATGAL+ NSCs in *Catnb^del(ex3)^* mice (p<0.0001). A.U.= Arbitrary Units. n=3 (**F**) YFP, GFAP, Sox2 and Ki67 immunolabelling in the DG of Control and Catnb^del(ex3)^ mice 30 days after tamoxifen administration. Scale bars, 50μm. (**G, H**) Quantifications of the data shown in (**F**) of the total number of NSCs normalised to SGZ length (mm) (Control vs Catnb^del(ex3)^: 30.95 ± 0.86 vs 16.65 ± 1.26, **G**) and the proportion of Ki67+ NSCs (Control vs Catnb^del(ex3)^: 1.33 ± 0.33% vs 3.8 ± 0.58%, **H**). NSCs are lost from the DG 30 after stabilising β-catenin and their proliferation is increased. n=3 for Control, n=5 for Catnb^del(ex3)^. (**I**) YFP, GFAP, Sox2 and Ki67 immunolabelling in the DG of Control and Catnb^del(ex3)^ mice 10 days after tamoxifen administration. Scale bars, 50μm. (**J, K**) Quantifications of the data shown in (**I**) of the total number of NSCs normalised to the length of the SGZ (mm) (Control vs Catnb^del(ex3)^: 32.38 ± 4.83 vs 28.92 ± 4.72, **J**) and the proportion of Ki67+ NSCs (Control vs Catnb^del(ex3)^: 4.33 ± 1.45% vs 7 ± 1.73%, **K**). Neither the total number of NSCs nor their proliferation are affected 10 days after stabilising β-catenin. n=3. (**L**) Representative images of SGZ located (arrowheads) and displaced (arrows) NSCs (YFP+ GFAP+ Sox2+ NSCs) in the DG of Control and Catnb^del(ex3)^ mice 10 days after tamoxifen administration. Dashed lines mark the SGZ. Scale bars, 50μm. (**M, N**) Quantifications of the data shown in (**L**). SGZ located NSCs (arrowheads) retain their correct niche location with their cell body located in the SGZ and a radial process through the GCL. Displaced NSCs in Catnb^del(ex3)^ mice (arrows) were identified as YFP+ GFAP+ Sox2+ NSCs that are located more than 2 cell nuclei away from the SGZ. The total number of displaced NSCs normalised to the SGZ length (mm) (Control vs Catnb^del(ex3)^: 1.42 ± 0.32 vs 4.59 ± 2.89, **M**) and the proportion of displaced NSCs (Control vs Catnb^del(ex3)^: 4 ± 0.58% vs 12.67 ± 6.57%, **N)**. Displacement of NSCs is increased in Catnb^del(ex3)^ mice compared with *Control*. n=3. SGZ, subgranular zone. GCL, granule cell layer. ML, molecular layer. Statistics: unpaired Student’s t-test (**D, E, G, H, J, K, M** and **N**). (ns, p>0.05. *, p<0.05. ***, p<0.001). Error bars represent mean with SEM.

**Figure S3:**
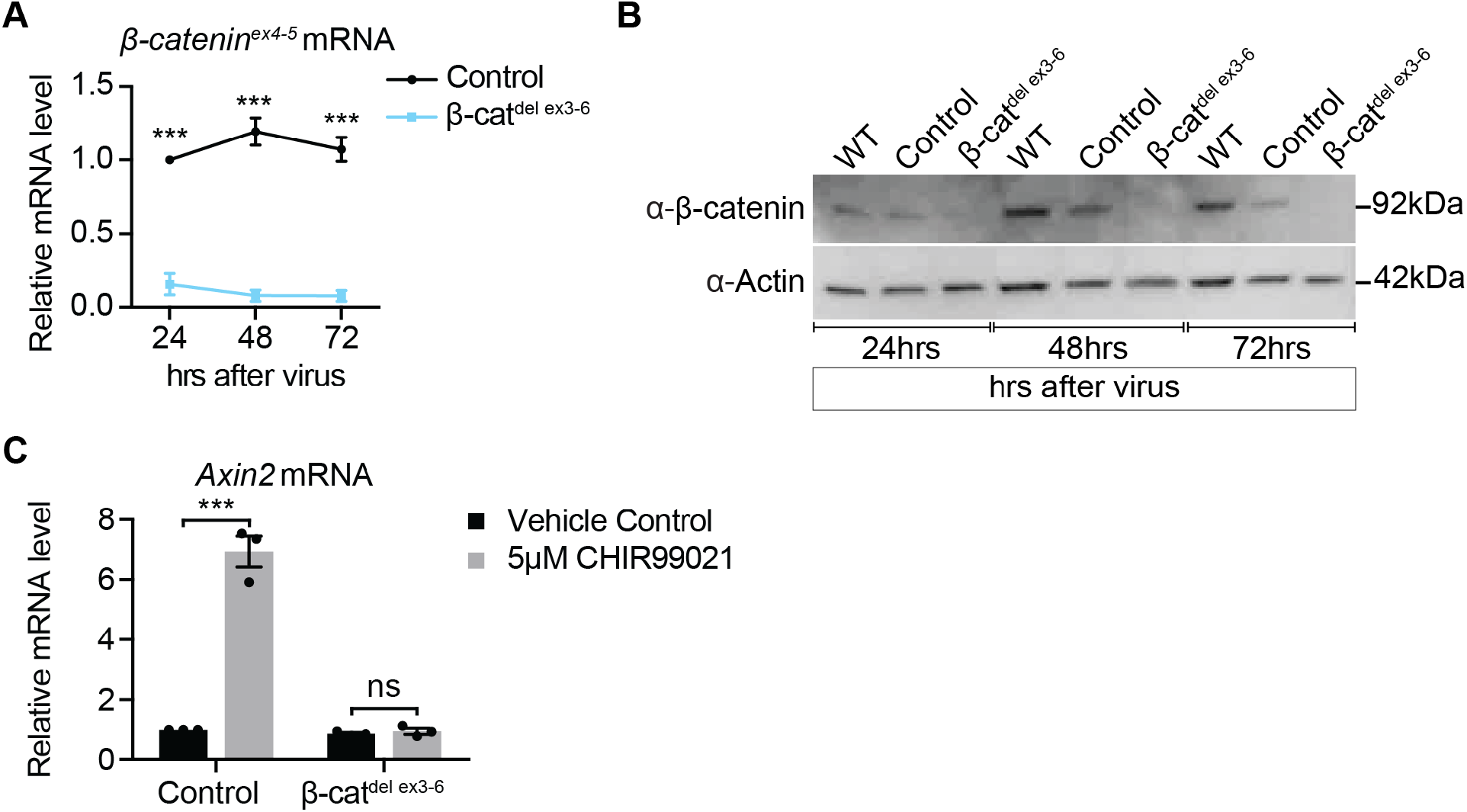
Validation of β-catenin recombination and deletion in β-cat^del ex3-6^ NSCs. (**A**) qPCR gene expression analysis of *β-cat^ex4-5^* (blue box, Figure S2**A**) 24hrs, 48hrs and 72hrs after control- and Cre-adenovirus transduction in Control and β-cat^del ex3-6^ active NSCs. *β-cat^ex4-5^* lies within the floxed region of the *β-cat^fl/fl ex3-6^* allele and is downregulated in active β-cat^del ex3-6^ NSCs vs Control (p<0.0001 for all time points). n=3. (**B**) Western blot of β-catenin protein levels in wild type (WT, no virus), Control (control-adenovirus) and β-cat^del ex3-6^ (Cre-adenovirus) active NSCs 24hrs, 48hrs and 72hrs after adenovirus transduction. β-catenin protein levels are abolished 48hrs after adenovirus transduction. n=3. (**C**) β-cat^del ex3-6^ active NSCs fail to upregulate *Axin2* in response to 5μM CHIR99021 treatment, indicating their inability to respond to a Wnt/β-catenin stimulus. CHIR99021 treatment began 48hrs after adenovirus transduction and maintained for 48hrs. n=3. Statistics: Two-way ANOVA with Sidak’s multiple comparisons test (**A, C**). (ns, p>0.05. ***, p<0.001). Error bars represent mean with SEM.

**Figure S4:**
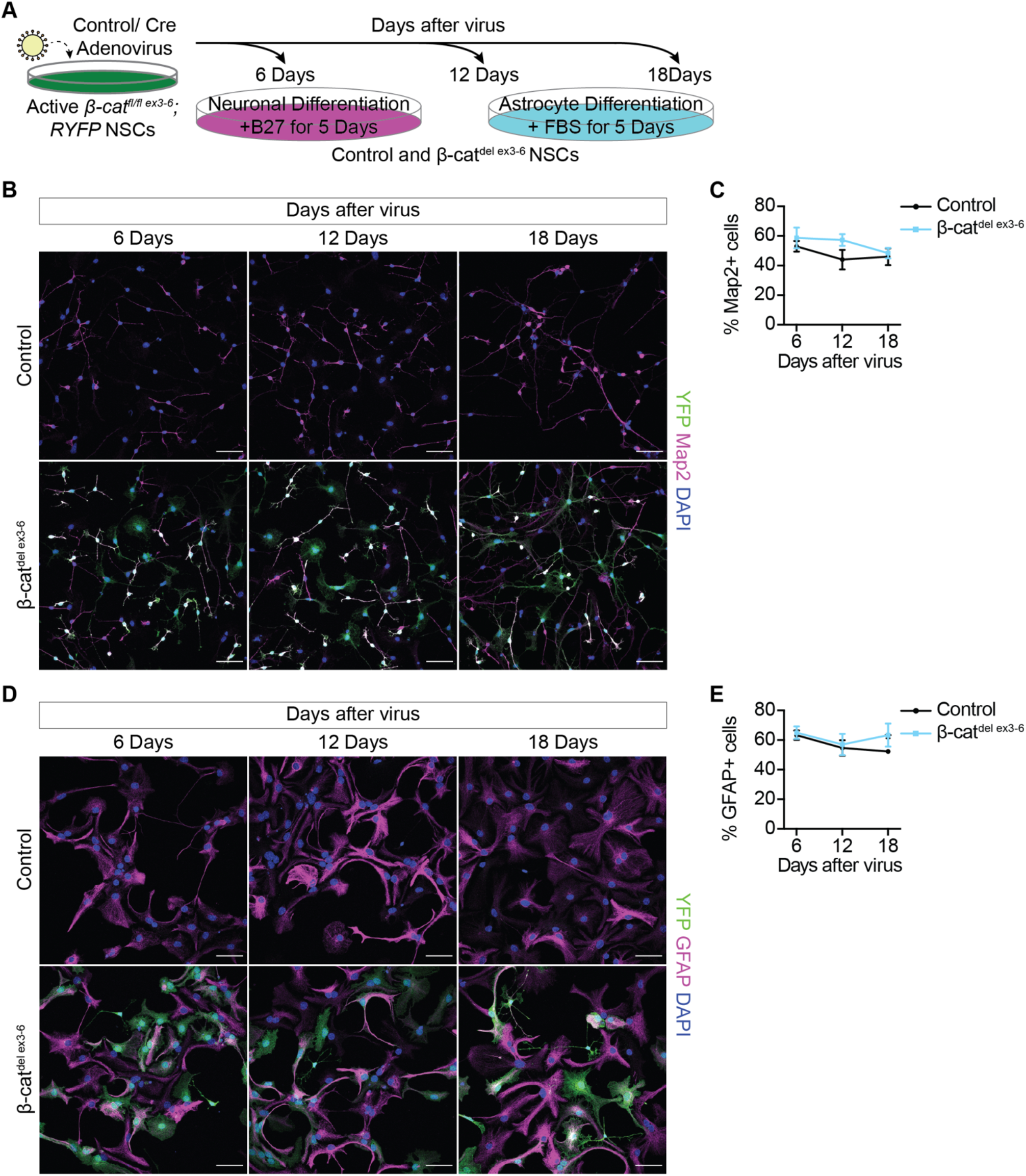
Chronic loss of Wnt/β-catenin signalling does not impair neuronal or astrocytic differentiation *in vitro.* (**A**) Six-, 12- and 18-days after virus transduction Control and β-cat^fl/fl ex3-6^ NSCs were cultured for 5 days in conditions to promote neuronal (+B27) and astrocytic (+FBS) differentiation. (**B**) YFP, Map2 and DAPI immunolabelling in Control and β-cat^del ex3-6^ NSCs cultured in B27 for 5 days at 6-, 12- and 18-days after virus transduction. YFP labels recombined cells and Map2 labels differentiated neurons. Scale bars, 50μm. (**C**) Quantification of the proportion of Map2+neurons in (**B**) (Control vs β-cat^del ex3-6^: 6days = 53 ± 3.61% vs 58.67 ± 6.96%, 12days = 44 ± 6.66% vs 57.33 ± 3.93%, 18days = 46 ± 5.69% vs 48.33 ± 3.53%). Control and β-cat^del ex3-6^ NSCs are similarly able to differentiate into neurons. Data shown are technical replicates of n=1 with an average of 147 cells counted per sample. (**D**) YFP, GFAP and DAPI immunolabelling in Control and β-cat^del ex3-6^ NSCs cultured in FBS for 5 days following 6-, 12- and 18-days after virus transduction. YFP labels recombined cells and GFAP labels differentiated astrocytes. Scale bars, 50μm. (**E**) Quantification of the proportion of GFAP+ astrocytes in (**D**) (Control vs β-cat^del ex3-6^: 6days = 63.33 ± 3.18% vs 65 ± 4.16%, 12days = 54.67 ± 5.18% vs 57 ± 7.21%, 18days = 52.33 ± 0.67% vs 63.33 ± 7.79%). Astrocytic differentiation is unaffected by loss of β-catenin. Data shown are technical replicates of n=1 with an average of 85 cells counted per sample. Error bars represent mean with SEM.

**Figure S5:**
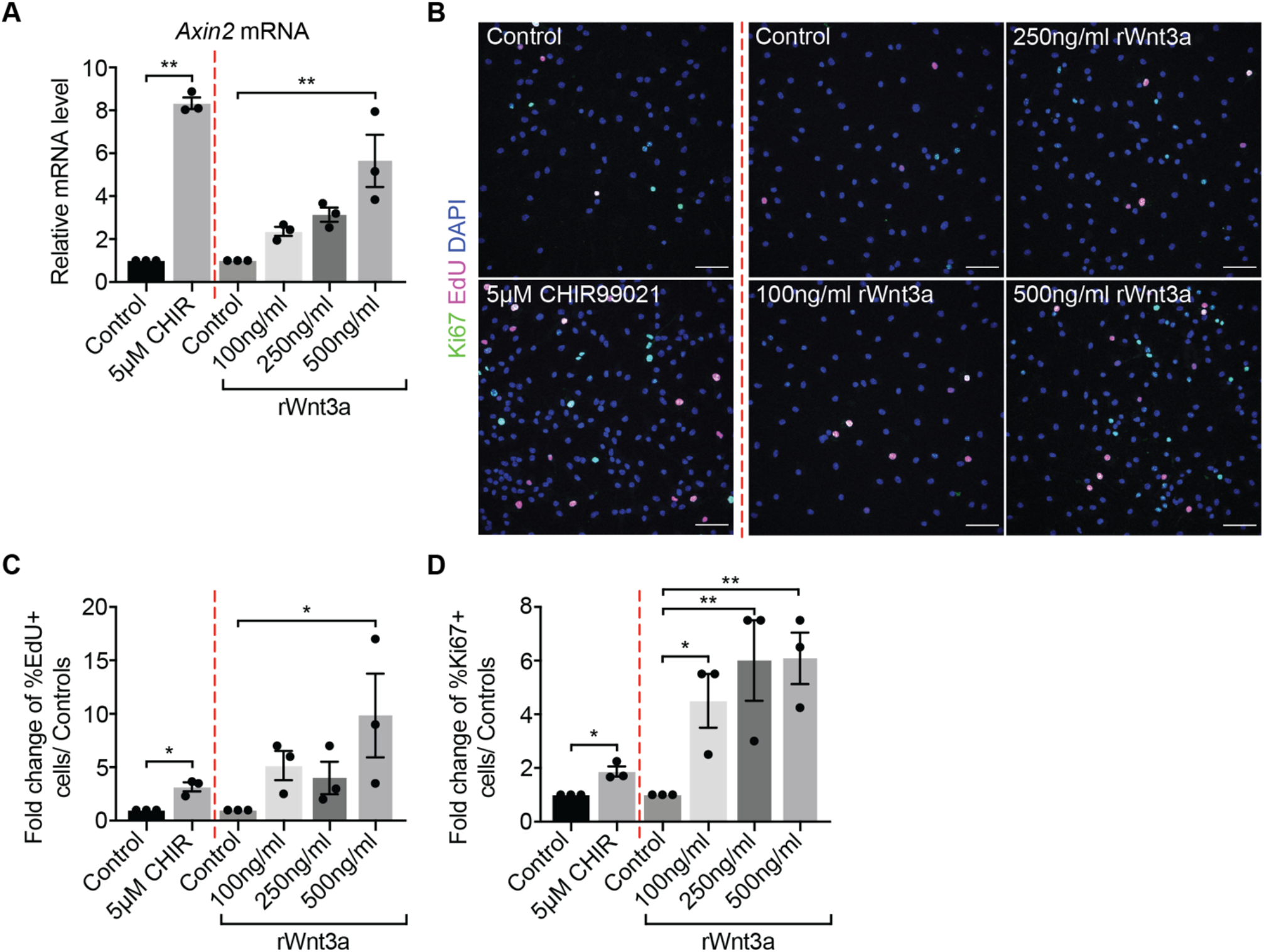
Treatment of quiescent NSCs with recombinant Wnt3a recapitulates the activation phenotype observed with CHIR99021. (**A**) *Axin2* expression is significantly upregulated by 48hrs sustained 5μM CHIR99021 and 500ng/ml Wnt3a treatment of quiescent NSCs. Results are shown normalised to the vehicle control for each Wnt/β-catenin agonist. n=3. (**B**) Immunolabelling of EdU, Ki67 and DAPI in quiescent NSCs treated with CHIR99021, rWnt3a and the vehicle controls for each agonist. Scale bars, 50μm. (**C, D**) Quantifications of the data shown in (**B**) of the fold change of proportion of EdU+ cells (**C**) and proportion of Ki67+ cells (**D**) for CHIR99021 and rWnt3a treated quiescent NSCs shown normalised to the vehicle control for each Wnt/β-catenin agonist. Both 5μM CHIR99021 and Wnt3a treatments increase proliferation of quiescent NSCs. n=3. Statistics: paired Student’s t-test for 5μM CHIR99021 compared with control and repeated measures one-way ANOVA with Dunnett’s multiple comparisons test for rWnt3a treatments compared with control (**A, C, D**) (ns, p>0.05. *, p<0.05. **, p<0.01). Error bars represent mean with SEM.

